# Automated high-throughput proteome and phosphoproteome analysis using paramagnetic bead technology

**DOI:** 10.1101/647784

**Authors:** Mario Leutert, Ricard A. Rodriguez-Mias, Noelle K. Fukuda, Judit Villén

## Abstract

Recent developments in proteomics have enabled signaling studies where >10,000 phosphosites can be routinely identified and quantified. Yet, current analyses are limited in throughput, reproducibility, and robustness, hampering experiments that involve multiple perturbations, such as those needed to map kinase-substrate relationships, capture pathway crosstalks, and network inference analysis. To address these challenges, we introduce rapid-robotic-phosphoproteomics (R2-P2), an end-to-end automated method that uses magnetic particles to process protein extracts to deliver mass spectrometry-ready phosphopeptides. R2-P2 is robust, versatile, high-throughput, and achieves higher sensitivity than classical protocols. To showcase the method, we applied it, in combination with data-independent acquisition mass spectrometry, to study signaling dynamics in the mitogen-activated protein kinase (MAPK) pathway in yeast. Our results reveal broad and specific signaling events along the mating, the high-osmolarity glycerol, and the invasive growth branches of the MAPK pathway, with robust phosphorylation of downstream regulatory proteins and transcription factors. Our method facilitates large-scale signaling studies involving hundreds of perturbations opening the door to systems-level studies aiming to capture signaling complexity.

## Introduction

Cellular signaling is organized as protein networks that respond to changing environments and diverse cellular needs to regulate cellular functions. Protein phosphorylation is integral to the signaling network, targeting most of the components: from sensors to kinases to effector proteins. To obtain a systems-level view of signaling it is crucial to obtain high dimensional measurements and deep coverage of the phosphoproteome. High dimensionality, obtained by sampling multiple perturbations and/or many timepoints after a perturbation, is required to define the edges of the network and obtain mechanistic insight into the structure and dynamics of the network. On the other hand, deep coverage is needed to quantify biologically relevant phosphorylation events that are low-stoichiometry and/or on low-abundance proteins (e.g. kinase activation loops).

Current phosphoproteomic methods allow identifying and quantifying thousands of phosphorylation sites. However, these methods still have limited reproducibility, robustness, and throughput, and therefore are not adequate for experiments involving tens to hundreds of perturbations. To facilitate systems biology studies of signaling we need a robust and automated end-to-end sample preparation workflow, along with mass spectrometry methods that provide systematic sampling at a discovery-scale.

Efficient end-to-end phosphoproteomic protocols have recently been developed to simplify, parallelize and, in some cases, automate sample preparation. Humphrey et al (Humphrey et al., 2015) have developed the EasyPhos platform, a streamlined manual approach that combines protein digestion, desalting and phosphopeptide enrichment starting from a cell lysate. In the newest version of the protocol sensitivity was improved by performing protein digestion and phosphopeptide enrichment in a single 96-well plate (Humphrey et al., 2018). Abelin et al (Abelin et al., 2016) have established a workflow for high throughput proteomic sample processing and phosphopeptide enrichment consisting of a robotic liquid handling platform (Agilent AssayMap Bravo) for automated protein reduction, alkylation and digestion, followed by a semi-automated peptide desalting step and an automated phosphopeptide enrichment over Fe^3+^-IMAC cartridges. These approaches allow deep and robust coverage of the phosphoproteome. However, as most commonly used proteomic sample preparation approaches, these methods rely on sample cleanup by solid-phase extraction on a reversed-phase C18 material. Therefore they are not compatible with many reagents (chaotropes, detergents, and polymers) commonly used for efficient cell lysis, challenging specimen, or subcellular organelle extractions. Inefficient removal of these reagents from the samples can inhibit enzymatic digestion and/or interfere with LC-MS/MS analysis. Instead, these methods use detergent alternatives, protein precipitation steps, phase-transfer protocols, extensive dilutions, molecular weight cutoff filters or affinity based methods leading to trade-offs in flexibility, sensitivity, throughput and handling (Jiang et al., 2004; Wiśniewski et al., 2009; Kulak et al., 2014).

An alternative approach to reversed-phase chromatography for universal proteomic sample preparation is a method called single-pot solid-phase enhanced sample preparation (SP3) that has recently been reported by Hughes et al. (Hughes et al., 2014), and later improved (Moggridge et al., 2018; Hughes et al., 2019). SP3 uses carboxylate-modified paramagnetic particles that capture proteins via aggregation induced by high organic solvents (Batth et al., 2019). The bead-protein complexes are washed to perform a protein cleanup followed by tryptic digestion in the same sample tube. The SP3 protocol is compatible with a wide variety of reagents (detergents, chaotropes, and salts) and allows elution of peptides in buffers that are directly compatible for LC-MS/MS analysis. Due to its high sensitivity, robustness and simple handling process SP3 has found broad application in low input proteomics (Hughes et al., 2014; Virant-Klun et al., 2016; Sielaff et al., 2017; Cagnetta et al., 2018; Buczak et al., 2018), however the implementation of SP3 in standard proteomic workflows has been limited, despite its advantages over reversed-phase chromatography methods and its obvious potential for automation and high□throughput sample processing.

Given the high performance and sensitivity of magnetic carboxylated microspheres for manual proteomic sample preparation we hypothesized that this methodology might be applicable to automated, high-throughput sample preparation using a magnetic particle processing robot. The flexibility of the method should furthermore allow for combination with downstream peptide enrichment technologies for post-translational modification proteomics. Tape et al (Tape et al., 2014) have shown that it is possible to use a magnetic particle processor in combination with magnetic microspheres to perform fully automated, highly reproducible phosphopeptide enrichment starting from a purified peptide mixture. In the current study, we systematically evaluated experimental parameters to implement an automated, high-throughput sample processing method based on paramagnetic beads that starts from cell lysates, performs protein capture, clean-up and digestion and is seamlessly combinable with automated phosphopeptide enrichment. We call our phosphoproteomic sample preparation method R2-P2 (rapid-robotic-phosphoproteomics), and the initial proteomics sample preparation R2-P1 (rapid-robotic-proteomics).

Reproducibility in phosphoproteomics should be extended beyond sample preparation and into the LC-MS/MS analysis. Most phosphoproteomics studies so far have employed data-dependent acquisition (DDA) MS measurements. DDA produces extensive data sets, however its stochastic sampling leaves many missing values when dealing with multiple samples. Data-independent acquisition (DIA) MS is a promising alternative for phosphoproteomics, achieving reproducible sampling, deep phosphoproteome coverage, good quantitative accuracy, and resolution of phosphopeptide positional isomers (Lawrence et al., 2016; Searle et al., 2018a).

We demonstrate the ability of R2-P2 in combination with DIA-MS to accelerate and leverage the quantitative analysis of phosphorylation sites in systems-biological studies involving multiple perturbations. Specifically, we study the phosphorylation dynamics of the mating, the high osmolarity and the invasive growth mitogen-activated protein kinase (MAPK) pathways, which share many signaling components, but result in very distinct cellular responses. We quantitatively measured the phosphoproteome of *S. cerevisiae* exposed to six different MAPK pathway related stimuli in a three point time course. We characterized global changes in signaling as well as pathway specific phosphorylation patterns.

## Results

### An automated magnetic sample preparation method for phosphoproteomics

We aimed at implementing a method for automated, high-throughput sample preparation using carboxylated microspheres on a magnetic particle processing robot, that could be seamlessly combined with automated phosphopeptide enrichment on the same robot. For this we designed the R2-P2 workflow that is conceptually based on the SP3 methodology from Hughes et al (Hughes et al., 2019), but executable in a 96-well format by a magnetic particle processing robot (KingFisher™ Flex). We configured the platform to perform R2-P1 in a first run and phosphopeptide enrichment for R2-P2 in a second run (Fig 1A). Briefly, the R2-P2 protocol starts by moving the carboxylated beads to a plate that contains protein extracts (e.g. cell lysates) and a high percentage of organic solvent, which promotes protein aggregation and capture on the beads. The carboxylated bead-protein complexes are subsequently passed through three individual plates containing high-organic solvents to wash the bead-protein complexes and remove salts, detergents, lipids, and other contaminants. The clean bead-protein complexes are then moved to a plate containing digestion enzyme in a buffered aqueous solution and the proteins are digested on the beads at 37**°** C and simultaneously eluted. We configured the binding and desalting steps to take 30 minutes and the digestion step 3.5 hours. At this point aliquots for total proteome analysis can be removed, dried and are ready for LC-MS/MS measurement. With the rest of the plate robotic phosphopeptide enrichment using Fe^3+^-IMAC, Ti^4+^-IMAC, Zr^4+^-IMAC or TiO_2_ magnetic spheres can be performed. For this we established a protocol based on previously established methods (Ficarro et al., 2009; Tape et al., 2014), which takes another 50 minutes and results in phosphopeptides that are ready for LC-MS/MS measurement after drying. The R2-P1 and R2-P2 protocols (SOP) and programs (*.bdz) are provided as supplementary files and can be run on any King Fisher™ Flex system.

**Figure 1.**
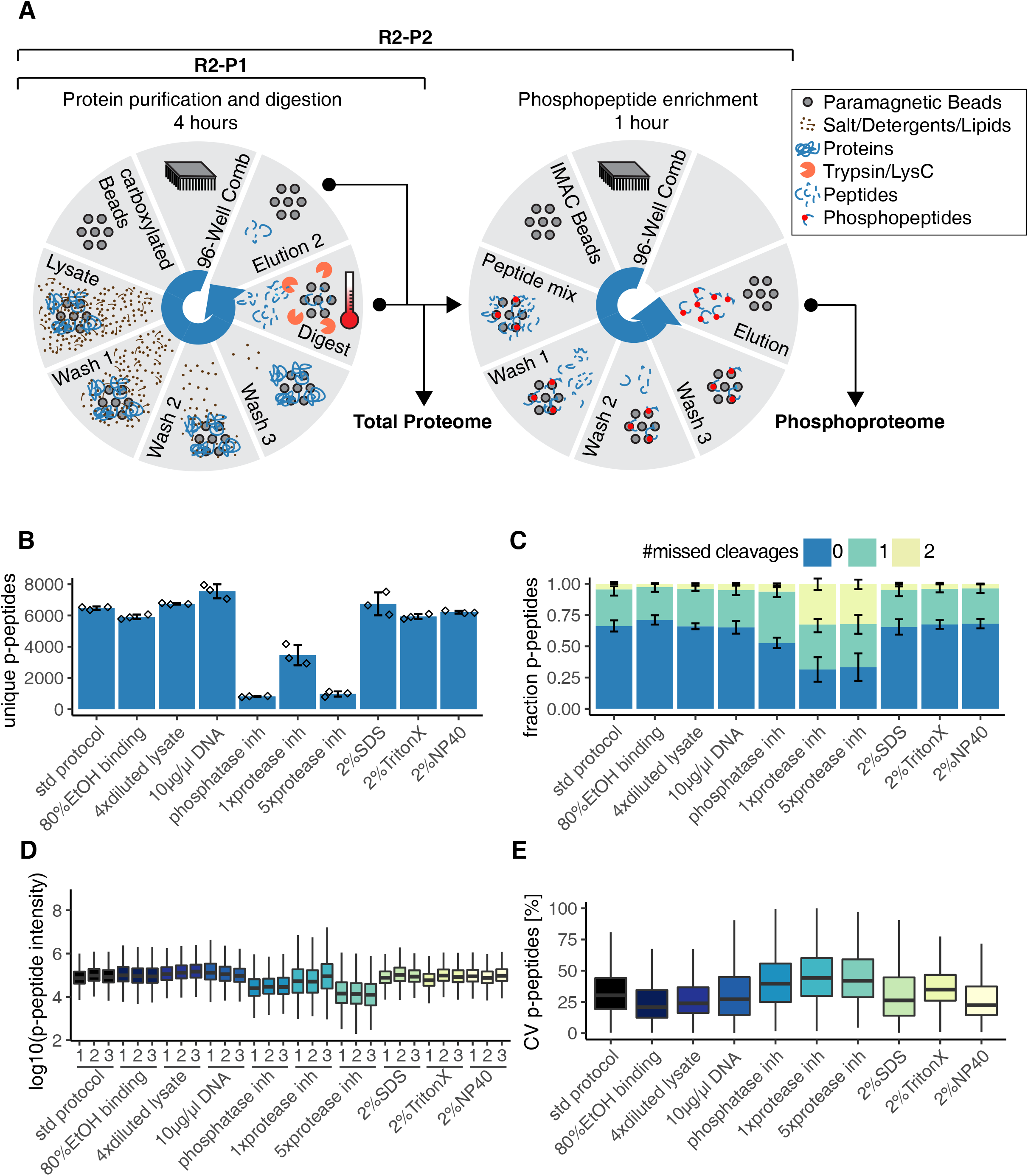
The R2-P2 method for high throughput robotic magnetic sample preparation for phosphoproteomic profiling. A) KingFisher^TM^ Flex configuration for R2-P2. The robotic configuration allows for loading of 8 different 96-well plates. Each plate can be rotated into position under a 96-pin magnetic head that drops down inside the 96-well plate to release, bind or agitate the magnetic microspheres in solution. In the first robotic run proteins are captured from lysates by carboxylated magnetic beads, purified, and eluted by digestion at 37°C. Eluted peptides are dried down and can be resuspended for total proteome analysis by LC-MS/MS and/or for automatic phosphopeptide enrichment. Phosphopeptides are enriched using a second robotic run on the KingFisher^TM^ Flex, using Fe^3+^-IMAC, Ti^4+^-IMAC, Zr^4+^-IMAC or TiO_2_ magnetic microspheres, and analyzed by LC-MS/MS to obtain the phosphoproteome. B) 200 µg of yeast protein extract in different buffer compositions was processed by R2-P2 followed by DDA-MS peptide analysis. Sample preparation was performed in triplicates. Mean count of unique phosphopeptides are shown for the different conditions (n=3). C) Fraction of phosphopeptides with 0, 1 or 2 missed cleavage sites. The mean is displayed (n=3). D) Boxplot depicting phosphopeptide MS1 signal intensity distributions. E) Boxplot depicting coefficients of variations (CVs) for replicate analysis of phosphopeptide MS1 signal intensities (n=3).

### R2-P1 allows efficient protein capture, desalting and peptide recovery

To establish an automated workflow for protein buffer exchange, reduction and alkylation, desalting, and on-bead digestion, we optimized conditions and assessed the efficiency of the following steps of the protocol: (1) protein binding to carboxylated beads, (2) peptide recovery from carboxylated beads and (3) digestion efficiency on carboxylated beads using different proteases.

In the original SP3 protocol, binding of proteins to carboxylated beads is induced by acetonitrile (50% v/v in lysate) under acidic conditions (Hughes et al., 2014). It was later reported that the acidic conditions reduce protein binding (Sielaff et al., 2017) and systematic study of a wide range of binding conditions proposed that ethanol at neutral conditions provided the best results in terms of binding, recovery and ease of use (Moggridge et al., 2018). To assess the efficiency of protein capture by R2-P1 we adjusted a yeast lysate to 50% or 80% acetonitrile or ethanol respectively (v/v in lysates) at pH 2 or pH 8. Binding at 50% or 80% ethanol at pH 8 and washing the beads with 80% ethanol provided the best results in terms of protein recovery (Fig S1A) and magnetic bead capture efficiency by the robot. This is in agreement with the newest SP3 protocol by Hughes et al (Hughes et al., 2019). For all further experiments protein capture at 50% ethanol at pH 8 and washes in 80% ethanol was used.

Digestion of proteins on beads in aqueous solution should lead to the elution of peptides from the beads, however it has been observed that this is not always efficient (Batth et al., 2019). We aimed at testing different elution steps for recovering digested peptides from the carboxylated beads during R2-P1. After digestion in 25 mM ammonium bicarbonate buffer (elution 1), beads were transferred to water and agitated for 5 min (elution 2) and then transferred to a 2% SDS solution, heated to 95ºC and sonicated (elution 3). The remaining proteins in the lysate, after R2-P1 protein capture, were separately digested (flow through). All fractions were analyzed by LC-MS/MS. The total median peptide intensity in elution 2 was 10% of elution 1, elution 3 was 6% of elution 1 and flow through was 12% of elution 1 (Fig S1B). The total ion current for the different elution steps confirmed that most of the protein mass is captured by the beads and peptides are efficiently eluted during the first elution/digestion step (Fig S1C). Analysis of the physicochemical properties of the proteins remaining in the flow through of the protein capturing step showed that R2-P1 is slightly biased against small (<30 kDa) and acidic proteins (Fig S1C). Based on the proposed mechanism of protein capture by the beads (Batth et al., 2019) it is likely that these proteins are not efficiently aggregated on the beads due to their solubility in organic solvents.

Next, we benchmarked different digestion conditions for R2-P1. A total yeast lysate was digested for 3.5 hour with either trypsin or LysC at an enzyme to protein substrate ratio ranging from 1:50 to 1:400. Both digestion enzymes provided good and comparable results up to a ratio of 1:200 as judged by total number of peptides identified (trypsin: n=13,198±144, Lys-C: n=12,868±180, Fig S1E) and percentage of fully cleaved peptides (trypsin: 70%, Lys-C: 86%, Fig S1F). We set on trypsin at 1:100 as our preferred enzyme for subsequent experiments.

### R2-P2 efficiently enriches phosphopeptides and is robust against a wide range of lysis buffers

We implemented R2-P2 on the same robot by coupling R2-P1 to automated Fe^3+^-IMAC phosphopeptide enrichment (Fig 1A). In a first step R2-P2 was performed on 200 µg yeast lysate, where proteins were captured either at 50% ethanol (std protocol) or at 80% ethanol (v/v in lysates). In both cases R2-P2 was able to efficiently enrich phosphopeptides with slightly more identifications for binding in the presence of 50% ethanol (n = 6467±104) versus 80% ethanol (n=5906 ± 146) (Fig1B).

Next we systematically evaluated lysis buffer composition compatibility for R2-P2. The following variations on our standard lysis buffer (8 M urea, 150 mM NaCl, 100 mM Tris pH 8) were tested: four times dilution of the lysis buffer by water; spike in of 10 µl/µg activated salmon sperm DNA; complementation with phosphatase inhibitors (50 mM sodium fluoride, 10 mM sodium pyrophosphate, 50 mM beta-glycerophosphate, 1 mM sodium orthovanadate); protease inhibitor mix (Pierce); 2% SDS; 2% NP-40 and 2% Triton-X. Samples were analyzed by LC-MS/MS before and after phosphopeptide enrichment. Total proteome analysis showed comparable peptide identifications (n~12,000) and digestion efficiencies for most conditions (Fig S2A, Fig S2B). Samples with protease inhibitors were an exception, showing a protease inhibitor concentration dependent decrease in peptide identification (Fig S2C) along with increased frequency of missed cleavage sites both with trypsin and Lys-C (Fig S2D). This suggests that certain compounds of the protease inhibitor mix are bound by the carboxylated beads and significantly inhibit enzymatic protein digestion. Of the three detergents tested only 2% SDS had a slight negative effect on peptide identifications (n=9,291±313), possibly because it was not efficiently removed during the 3 washes and interfered with LC-MS/MS analysis. If such high SDS concentrations are to be used, an additional carboxylated bead washing step may improve the results.

Starting from 200 µg yeast lysate, R2-P2 showed comparable phosphopeptide identifications for the different lysis buffer compositions (n~6,000) with drastically reduced identifications for the lysis buffer containing phosphatase inhibitors (n = 814±40), 5x protease inhibitor mix (n = 975±178) or 1x protease inhibitor mix (n = 3,460±644) (Fig 1B). In line with the results from total proteome analysis, we observed that a high percentage of the phosphopeptides identified contained missed cleavage sites in samples with 5x and 1x protease inhibitor mix (67% and 69% missed cleaved peptides). This percentage was lower for samples with phosphatase inhibitors (48% missed cleaved) and down to the expected levels in all other conditions (< 35% missed cleaved) (Fig 1B). To assess how these conditions perform on a quantitative experiment we measured MS1 phosphopeptide precursor intensities. All conditions showed comparable values averaging 10^5^ intensity units, except for the samples containing phosphatase or protease inhibitors, which were an order of magnitude lower (Fig 1C). Quantitative reproducibility assessed by the distribution of coefficients of variation (CV) was as good as is expected for label-free quantification experiments (CV < 30%) and again, significantly worse for protease or phosphatase inhibitor containing samples (CV > 40%) (Fig 1D). We conclude that similar to the protease inhibitor binding effect described above, phosphatase inhibitors are also co-enriched by the carboxylated beads. These do not introduce problems for total proteome analysis, but are strongly impairing phosphopeptide enrichment, likely due to competition for Fe^3+^ binding.

Taken together these results show that the R2-P2 provides a robust method for high-throughput sample preparation for total proteome as well as phosphoproteome analysis that is compatible with a wide variety of detergents and chaotropic agents. Importantly, we identified severe problems in using protease and/or phosphatase inhibitors in the lysis buffer. In lieu of these inhibitors, we recommend blocking endogenous enzymatic activities by using harsh, denaturing lysis buffers.

### Benchmarking and scalability of automated magnetic sample preparation

To benchmark our method we compared its performance to the widely used method of preparing proteomic and phosphoproteomic samples, which involves in-solution digestion and desalting by solid-phase extraction on C18 SepPak cartridges. First, we processed 25 µg yeast protein extract for total proteome analysis by the two methods. Analysis of ~0.5µg by LC-MS/MS revealed more peptide identification by R2-P1 (n = 13,237 ± 476) than by SepPak (n = 11,011 ± 323) (Fig S3A). Comparison of all identified peptides showed that for R2-P1 74% of peptides were identified in at least two replicates and for SepPak 63%. While around 50% of all peptides were identified by both methods (Fig S3B), protein coverage was better with R2-P1 (Fig S3C). Median MS1 intensity of the identified peptides was slightly higher in the SepPak method (median log_10_ (intensity) = 6.96) compared to R2-P1 (median log_10_(intensity) = 6.67) (Fig S3D), however median CVs for the identified proteins were better for R2-P1 (CV = 20.4%) versus SepPak (CV = 24.6%) (Fig S3E). Comparison of molecular weights, GRAVY scores and isoelectric points for the identified peptides (Fig S3F) and corresponding proteins (Fig S3G) showed again a slight bias of the magnetic bead method against small acidic proteins.

To compare performance, scalability and optimal protein starting material for the two methods coupled to automatic phosphopeptide enrichment we processed 25, 50, 100, 200 and 400 µg of a yeast lysate in parallel and injected one fourth of the enrichment into the MS. For both methods, the number of identified phosphopeptides scaled according to the starting protein amount reaching a maximum number of phosphopeptide identifications at 200 µg for R2-P2 (n=6,435 ± 301) and at 400 µg for SepPak (n=5,758 ± 542) (Fig 2A). With 25 µg of starting material, R2-P2 was still able to identify a high number of phosphopeptides (4,176 ± 619) that was significantly higher compared to SepPak identifications (2,904 ± 217), confirming the suitability of solid-phase enhanced sample preparation for low input proteomics. As expected, phosphopeptide intensities increased up to 400 µg for both methods (Fig 2B). Replicate comparison showed that median CVs for phosphopeptides identified by R2-P2 ranged from 25.3% (400 µg) to 33% (50 µg), whereas median CVs for SepPak processed samples ranged from 25.7% (400 µg) to 40% (200 µg) (Fig 2C). Overlap between all identified phosphopeptides by the SepPak based method and R2-P2 for 3 individually processed and singly injected samples was between 45% (25 µg starting amount) and 60% (100 µg starting amount) (Fig 2D), which was comparable to the overlap between two replicates of either method. Analysis of the 200 µg sample showed that R2-P2 was covering more phosphosites per protein (on average 3.5 versus 3 sites per protein) (Fig 2E).

**Figure 2.**
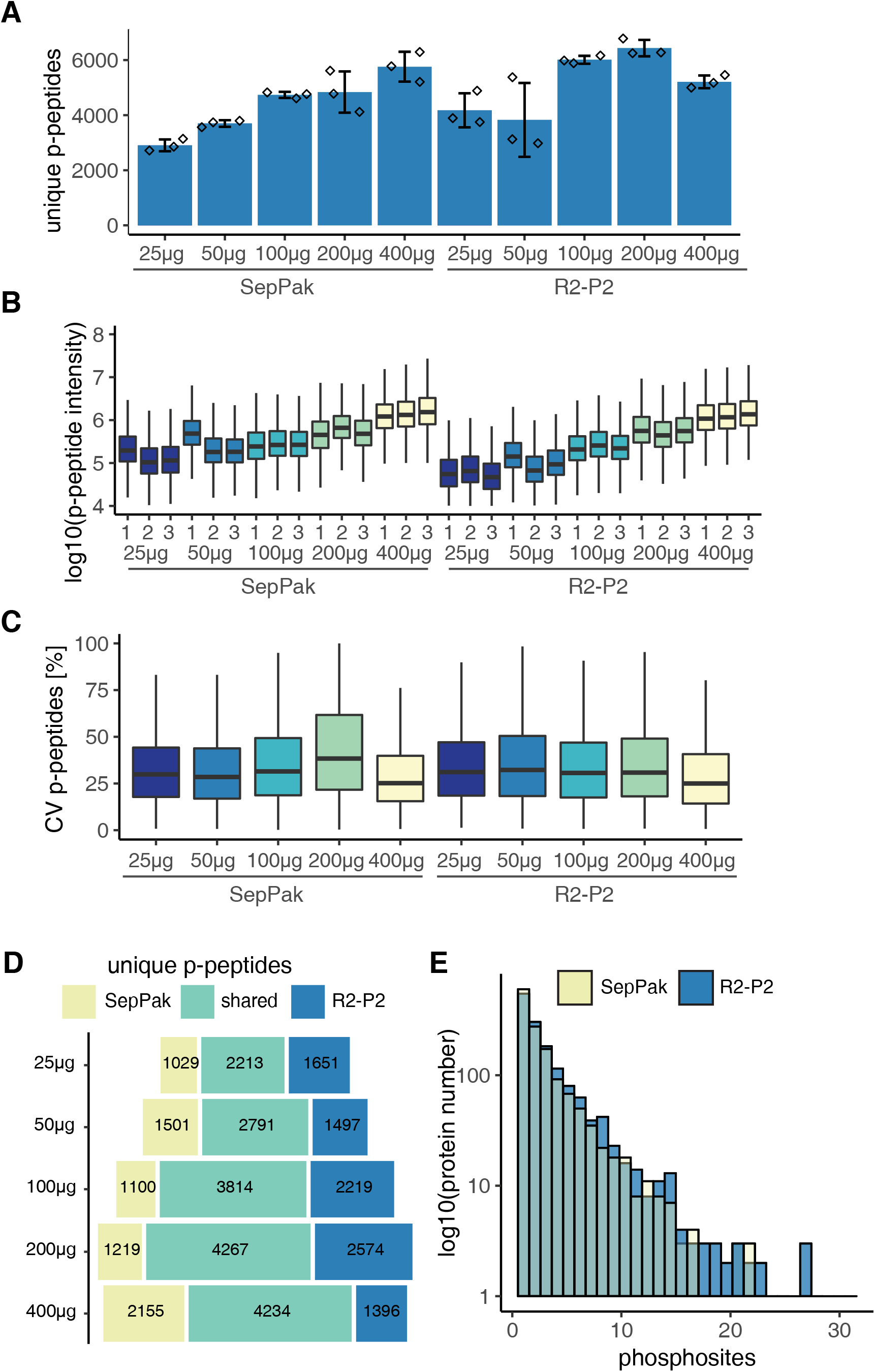
R2-P2 allows scalable, high sensitivity phosphoproteomics. Different amounts (25 µg, 50 µg, 100 µg, 200 µg and 400 µg) of yeast protein extract were processed by R2-P2 or by manual in-solution digestion and reversed phase C18 desalting (SepPak), phospho-enriched by robotic Fe^3+^-IMAC followed by peptide analysis with DDA-MS. Sample preparation was performed in triplicates. A) Mean count of unique phosphopeptides are shown (n=3). B) Boxplot depicting phosphopeptide MS1 signal intensity distributions. C) Boxplot depicting CVs for replicate analysis of phosphopeptide MS1 signal intensities (n=3). D) Overlap of all identified phosphopeptides. E) Histogram analysis of the number of phosphosites per protein detected by the different methods for 200 µg protein input.

Taken together these results show that R2-P2 for total proteome and phosphoproteome analysis is directly comparable to in-solution digestion and C18 SepPak purification, the most commonly used proteomic sample preparation method. With R2-P2, we identified more unique phosphopeptides and achieved comparable or better reproducibility for label free quantification. Importantly, R2-P2 performs significantly better with low sample amounts than the SepPak method. Phosphopeptide identifications maxed out at 100-200 µg and provided the most reproducible quantitative measurements at 400 µg. R2-P2 coupled to phosphopeptide enrichment is scalable and highly sensitive.

### Modularity of R2-P2 allows combination of different phosphopeptide enrichment methods

Given the modularity of our robotic approach, we sought to compare and benchmark the performance of different methods and materials for phosphopeptide enrichment in combination with R2-P2. For this, we conducted an experiment were triplicates of 250 µg of a common yeast protein extract were processed by R2-P2 and phosphopeptides were enriched using either Fe^3+^-IMAC, Ti^4+^-IMAC, Zr^4+^-IMAC or TiO_2_ microspheres. All materials performed well, highlighting the versatility of the method. We identified the highest number of phosphopeptides using Fe^3+^-IMAC (n=13,417 ± 368), followed by Ti^4+^-IMAC (n=12,254 ± 80), TiO_2_ (n=11,346 ± 1122), and Zr^4+^-IMAC (n=10,840 ± 852) (Fig 3A). Likewise, Fe^3+^-IMAC achieved the highest enrichment selectivity – on average 94% of all measured peptides were phosphopeptides – (Fig 3B) and recovered the highest fraction of multiple phosphorylated peptides (Fig 3C). Phosphopeptide precursor MS1 intensities were the highest with Fe^3+^-IMAC and TiO_2_ (Fig 3D) and all enrichments showed good reproducibility with median CVs around 25% (Fig 3E). Overlap of all identified phosphopeptides for the different enrichment methods showed significant portions that were exclusively identified by a single enrichment type (51% of all identified phosphopeptides) (Fig 3F).

**Figure 3.**
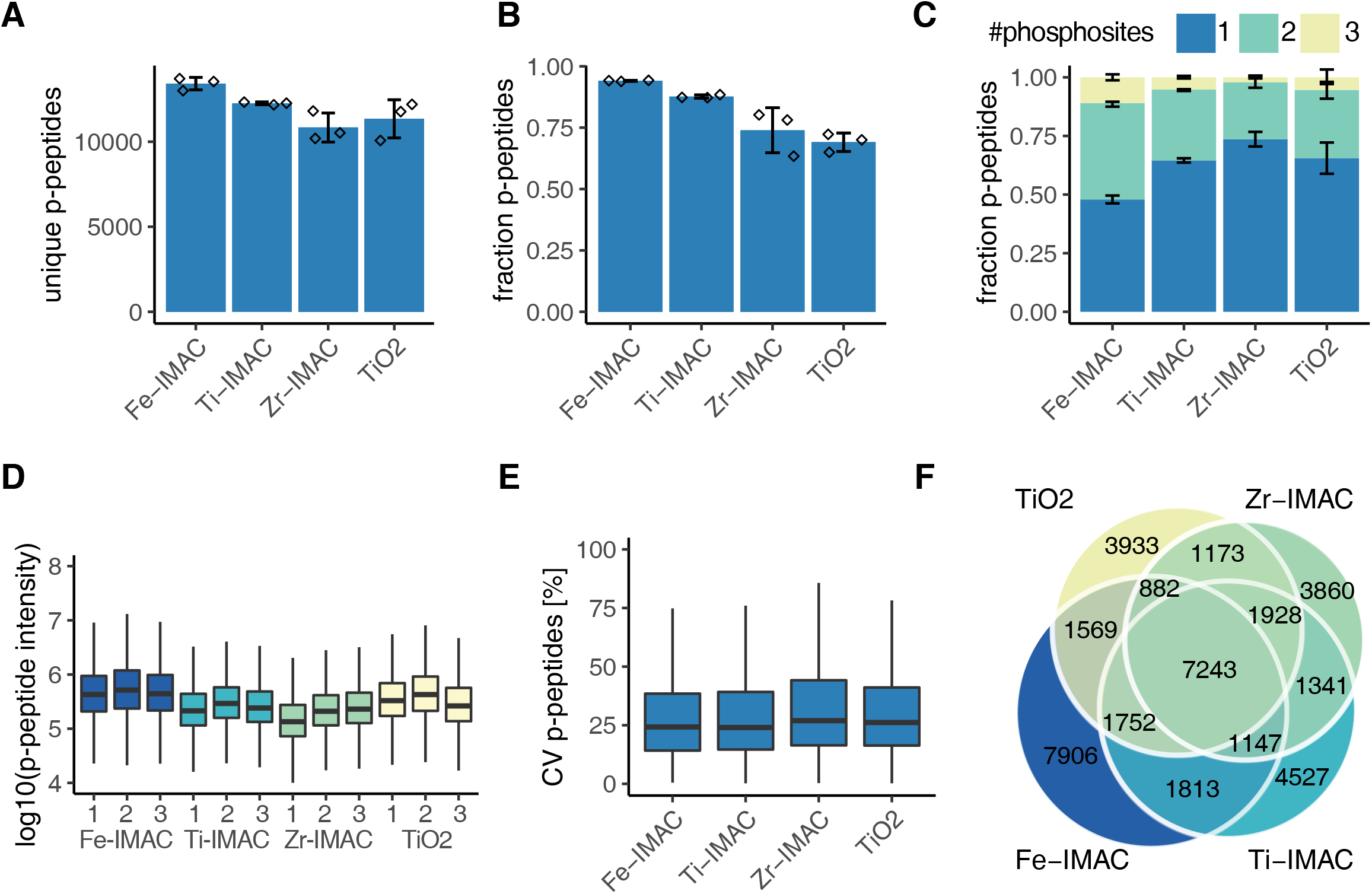
Combination of R2-P2 with different phosphopeptide enrichment methods allows for deep phosphoproteomic profiling. 250 µg of yeast protein extract was processed by R2-P2 using either Fe^3+^-IMAC, Ti^4+^-IMAC, Zr^4+^-IMAC or TiO_2_ magnetic beads and followed by peptide analysis with DDA-MS. Sample preparation was performed in triplicates. A) Mean count of unique phosphopeptides identified by the different enrichments are shown (n=3). B) Fraction of phosphorylated peptides versus unmodified peptides (n=3). C) Fraction of phosphopeptides with 1, 2 or 3 co-occurring phosphorylation sites (n=3). D) Boxplot depicting MS1 phosphopeptide signals. E) Boxplot depicting CVs for phosphopeptide MS1 signal distributions (n=3). F) Venn diagrams of all identified phosphopeptides by the different phospho enrichments

These results demonstrate the modularity and versatility of R2-P2 and highlight the benefits of using different methods and materials for phosphopeptide enrichment in combination, to obtain higher coverage of the phosphoproteome.

### Phosphoproteomics in a multiple perturbation experiment using R2-P2 and data independent acquisition mass spectrometry

We showed that R2-P2 allows the parallel and reproducible preparation of up to 96 samples for phosphoproteomic analysis. Here we combined R2-P2 with DIA-MS to systematically study cellular signaling along the MAPK axis, including the pheromone response/mating pathway, the high osmolarity stress response pathway and the invasive growth response pathway.

Experiments were performed in the Σ1278b *Saccharomyces cerevisiae* strain. Σ1278b has a fully functional invasive response pathway that can be induced by nutrient limitations and certain types of alcohols, whereas most laboratory yeast strains have acquired mutations that compromise the invasive growth response (Cullen and Sprague, 2012). Yeast cultures were exposed to one of 3 distinct stimuli (alpha factor, sodium chloride, and 1-butanol) or 3 media replacements (replacement of glucose with galactose, glucose limitation, and nitrogen limitation) or left untreated for 10, 30 and 90 minutes, in three biological replicates. Alpha factor induces the MAPK mating pathway and NaCl induces the MAPK high osmolarity glycerol (HOG) pathway. Replacement of glucose with galactose, glucose and nitrogen limitation, and 1-butanol have been described to activate the invasive growth pathway via MAPK and/or three other pathways (RAS/PKA, SNF and TOR) (Cullen and Sprague, 2012). For every sample, 400 µg yeast protein extract was processed using R2-P2. Given the orthogonal results from the different phosphopeptide enrichment methods, we decided to split the resulting peptide mixtures in two and enrich with Fe^3+^-IMAC and Ti^4+^-IMAC in parallel to obtain a deeper coverage of the phosphoproteome (Fig 4A).

**Figure 4.**
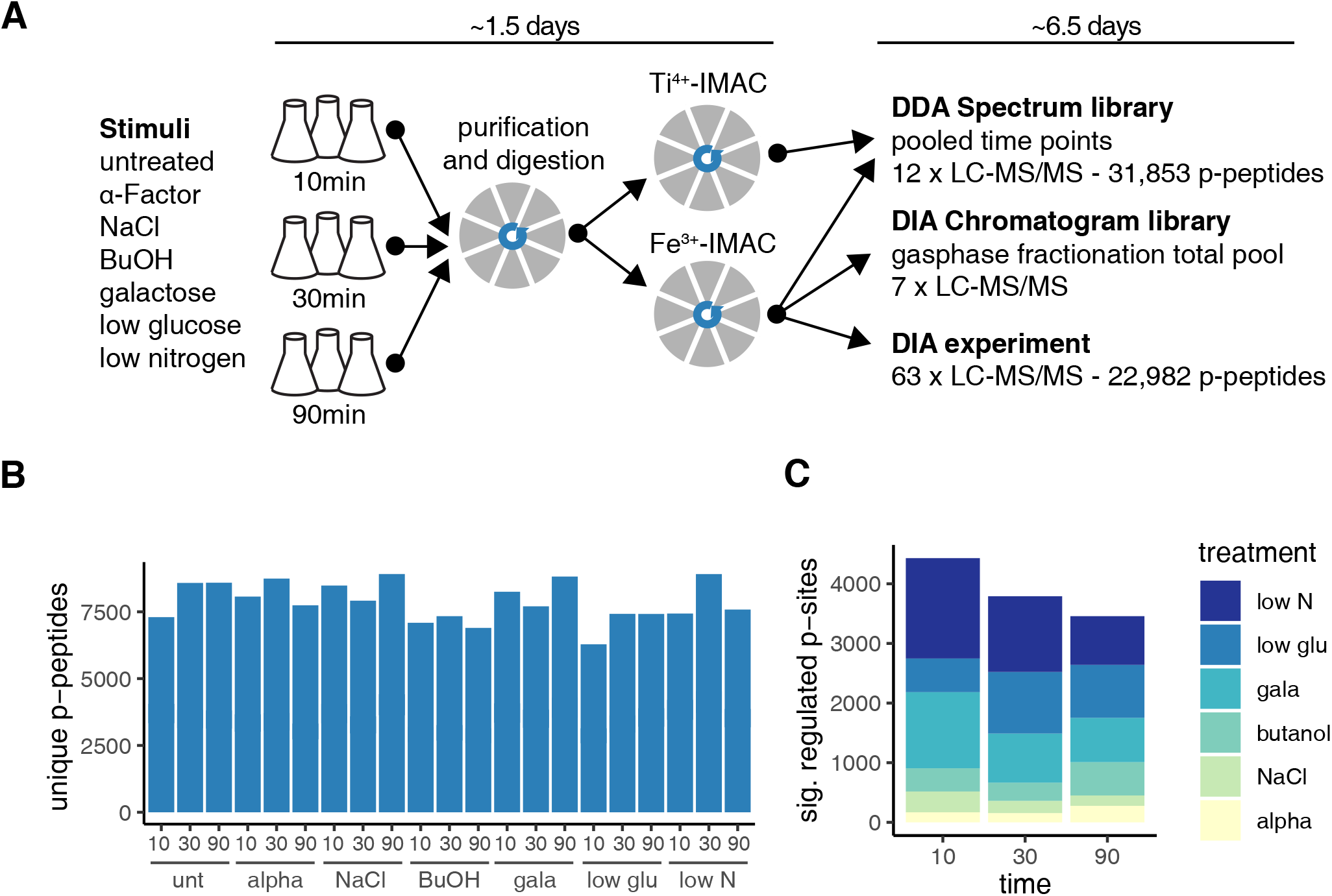
High-throughput, multi-condition phospho signaling mapping using R2-P2 and DIA-MS. A) Scheme of the experimental setup. Σ1278b MATa yeast cultures were grown and treated in biological triplicates with the indicated stimuli in a three point time course. Proteins were purified and digested using R2-P2 with Fe^3+^-IMAC or Ti^4+^-IMAC. The spectrum library was generated by DDA measurement of pooled timepoints of Fe^3+^-IMAC and Ti^4+^-IMAC enriched samples. The chromatogram library was generated by measuring a pooled sample from Fe^3+^-IMAC enrichment of all conditions and timepoints in seven staggered narrow-window DIA-MS runs. The Fe^3+^-IMAC fractions of each samples were measured in a single injection wide window DIA-MS experiment. The time used for sample preparation and measurement is indicated on the top and numbers of identified phosphopeptides on the right. B) Unique phosphopeptides present in at least two out of three replicates identified in the DIA-MS experiment. C) Significantly regulated phosphosites compared to the corresponding untreated control as determined by a two sample t-test (permutation based FDR < 0.01, and fold change > 2) for the different stimulation and time points.

For the LC-MS/MS measurement, we first acquired a limited number of DDA measurements to build a spectral library. For this, we pooled three biological replicates and the three time points for each condition and separately measured the Fe^3+^-IMAC and Ti^4+^-IMAC fractions. These 12 injections resulted in the measurement of 31,853 unique phosphopeptides (Fig S4A, Dataset EV1). Additionally, seven injections of a total sample pool were dedicated to gas-phase fractionation by staggered narrow-window DIA and were later used to generate a chromatogram library that catalogs retention time, precursor mass, peptide fragmentation patterns, and known interferences that identify each peptide within a specific sample matrix (Fig 4A) (Searle et al., 2018b). Subsequently, a single injection of the Fe^3+^-IMAC fraction for every condition (N = 63) was measured by wide window DIA for the label free quantitation experiment. The MS measurements resulted in 82 injections with 60 min separation gradients; together with the sample preparation (starting from cell pellets) the experiment was completed in 8 days (Fig 4A). For the analysis of the acquired data, we used Skyline to generate the spectral library from the DDA runs (MacLean et al., 2010), Encyclopedia to generate the chromatogram library from the narrow-window DIA runs (Searle et al., 2018b), and Thesaurus to identify, site-localize, and quantify phosphopeptides in the wide-window DIA runs (Searle et al., 2018a).

An average of 7,880 phosphorylated peptides were quantified in at least two out of three biological replicates for each condition and 40% of the phosphopeptides had all sites confidently localized (Fig 4B, Dataset EV2, Dataset EV3). Median CVs for the biological triplicates were between 13% to 33% with lower reproducibility for the 90 min time points, which likely reflects biological variability of phosphorylation in these samples (Fig S4B). Pearson’s correlation coefficients showed strong correlation between untreated, alpha factor, NaCl and BuOH treated samples and lower correlation between nutrient limitations (Fig S4C). Looking at localized phosphosites that were significantly changing in abundance compared to the untreated (two sample t-test, permutation based FDR < 0.01, fold change > 2) revealed that nutrient changes (i.e. low nitrogen, exchanging glucose for galactose, and low glucose) had the most profound impact on the phosphoproteome (Fig 4C).

### Global temporal dynamics of regulated phosphosites

To assess the similarity between the 19 conditions, we performed principal component analysis of the phosphoproteomics data (Fig S5A). Most samples separated well into groups according to the type and time of stimulation, supporting data reproducibility and outlining global correlation between stimuli (Fig S5A). Component 1 separates the nutrient limiting conditions from the untreated, alpha factor, NaCl and BuOH stimulated samples. Gene ontology enrichment analysis showed that separation in component 1 is driven by phosphoproteins involved in translation, signal transduction and kinase activity, as well as transmembrane transport, lipid and protein metabolic processes (Fig S5B). Component 2 resolves the temporal effect of the stimulation and is enriched in nucleic acid binding phosphoproteins and phosphoproteins with functions in RNA metabolic processes and transcription factors (Fig S5B). To gain more resolution in our analysis we conducted separate PCA analyses for each timepoint. Time-resolved PCA analysis showed good separation between different stimuli and close similarity between biological replicates for the 10 and 30 min time points (Fig 5A). The 90 min time point resulted in bigger spread in biological replicates, particularly for the osmotic stress with NaCl, suggesting they may follow different stress recovery trajectories (Fig 5A).

**Figure 5.**
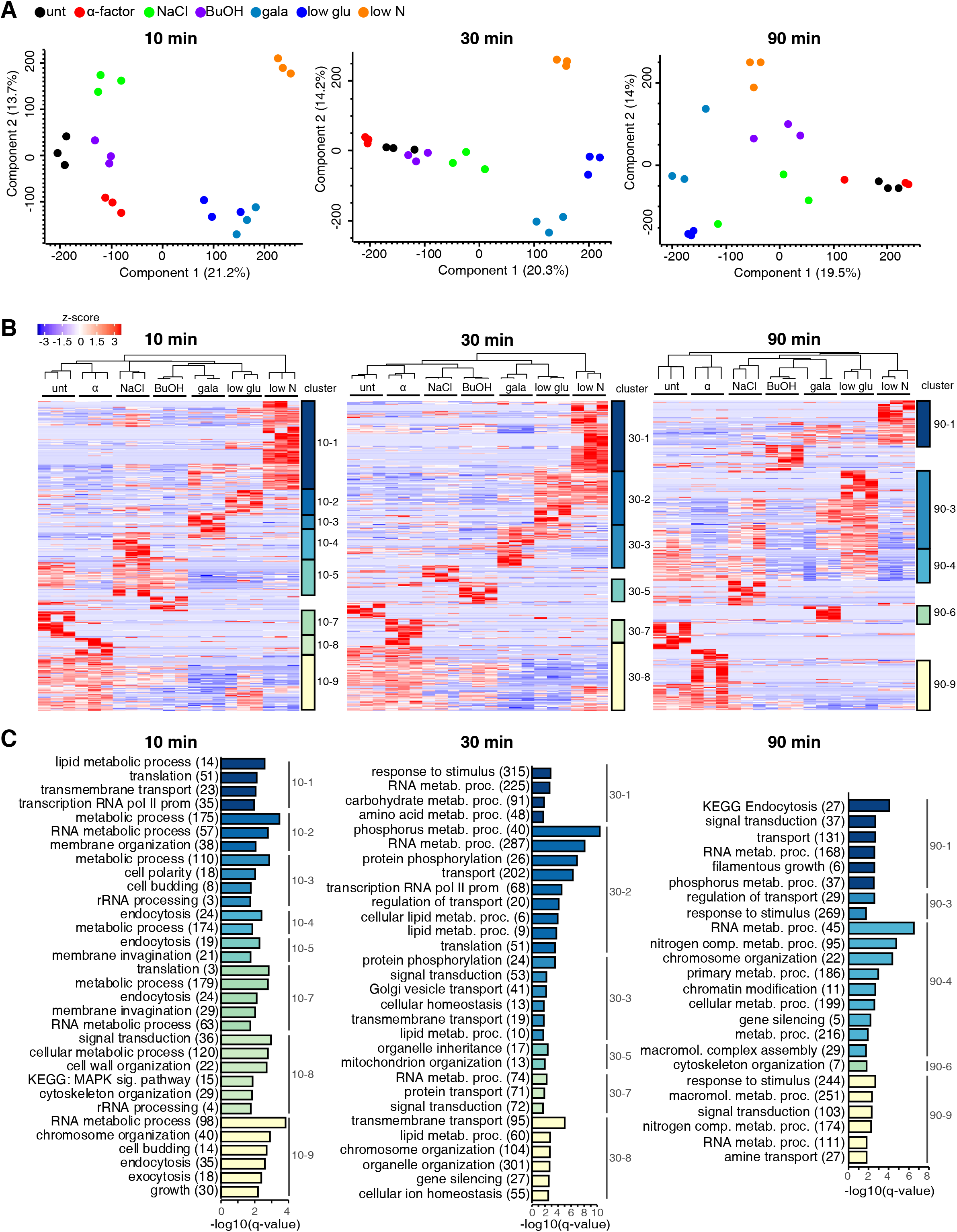
Analysis of time-resolved stimuli. A) Principal component analysis (PCA) of the DIA-MS phosphoproteomic data for the different stimuli separated by time points. B) Hierarchical clustering analysis of z-scored intensity values of significantly regulated (ANOVA, permutation-based FDR <0.05) phosphopeptides of the different time points. Column clustering hierarchy is indicated at the top. Row clusters for term enrichment analysis are color coded and indicated on the right. C) Term-enrichment analysis displaying gene ontology biological processes (GOBP slim) and Kyoto Encyclopedia of Genes and Genomes (KEGG) terms that were significantly enriched (Fisher’s exact test with Benjamini–Hochberg multiple-hypotheses correction at P < 0.02) for phosphorylated proteins in the corresponding clusters in B).

Next, we performed time-resolved hierarchical clustering of significantly regulated phosphopeptides (Fig 5B). Clustering of the 10 and 30 min time points showed similar characteristics with many of the clusters being selective to one condition, higher number of regulated phosphosites for the nutrient switch conditions and fewer for the salt stress. In contrast, 90 min clusters comprised phosphosites regulated in multiple conditions. We observed partially overlapping response for the nutrient switch conditions, with low glucose and low nitrogen overlapping at 30 min (cluster 30-2), and low glucose and galactose overlapping at 90 min (cluster 90-3). We also observed a sustained increase in the number of upregulated sites over time under low glucose conditions.

GO enrichment analysis of biological processes in the nitrogen starvation clusters identified a role for upregulated phosphoproteins in translation (10 min), in transcription and lipid metabolic processes (10 min and 30 min), invasive growth (90 min), and RNA metabolic processes and transmembrane transport (all timepoints). Phosphoproteins upregulated under low glucose or galactose are enriched in proteins involved in general metabolism, lipid metabolism processes and Golgi vesicle transport (30 min), and cytoskeletal organization (90 min). Butanol treatment only showed significantly enriched biological process GO terms at 30 min (cluster 30-5), which were organelle inheritance and mitochondrion organization. NaCl treatment showed the most pronounced treatment specific effect at 10 min (cluster 10-4). Similarly, the effect of alpha factor treatment was most pronounced at 10 and 30 min (cluster 10-8, 30-7) enriched in components of the MAPK signaling pathway, signal transduction, and organization of the cell wall and cytoskeleton, according to the induced mating response.

### Pathway analysis reveals stimulation-specific signatures on regulatory proteins and transcription factors

To identify activated kinases and pathways, we looked in our data for changes in phosphorylation-based sentinels, i.e. phosphopeptide markers that are indicative for the activity of pathways or biological processes (Soste et al., 2014). We found evidence for activation of Pka and Snf1 kinases in nitrogen and glucose limitation, galactose, and butanol treatment (Fig S6) indicative of activation of invasive growth. Given that developmental programs such as mating and invasive growth require alterations in cell cycle control, we expected cell cycle specific phosphopeptide markers to be regulated under the tested conditions. Indeed, alpha factor down regulated activity markers of the cell cycle kinases Cdc28 and Ck2 (Fig S6), indicative of G1 arrest to prepare for cell-cell fusion. We also found that Sch9 T723/S726, a sentinel of TORC1 activity, was suppressed under all nutrient limitations, and transiently reduced with NaCl (Fig S6, Fig S7), as previously reported (Urban et al., 2007).

To further investigate the effect of the treatments on TORC1 signaling, we mapped regulated phosphosites on a previously reported set of TORC1 components and effectors (Oliveira et al., 2015). We found multiple phosphosites on the TORC1 subunit Tco89 being regulated upon nitrogen and glucose starvation and to a lesser extent with galactose. Similarly, we found phosphosites on several regulatory proteins (Nap1, Aly2, Eap1, Igo1, Atg13) and transcription factors (Tod6, Dot6, Stb3, Fhl1, Ifh1, Maf1, Msn2, Msn4) downstream of TORC1 that were regulated under nutrient limitation (Fig S7). The transcriptional repressors Maf1, Tod6, Dot6 and Stb3 regulate ribosomal biogenesis and are directly controlled by inhibitory phosphorylation through the nitrogen-sensitive TORC1-Sch9 and the carbon-sensitive PKA axis, thereby adjusting protein synthesis to nutrient availability (Huber et al., 2011). As expected, we see canonical Sch9 sites selectively downregulated (Maf1 S90, Tod6 S280, Stb3 S285, Stb3 S286) upon low nitrogen (Huber et al., 2009, 2011) and upregulated under low glucose; but other sites on the same proteins showed opposite trends. This differential regulation can be explained by varying activity of the TORC1-Sch9 axis, the opposing TORC1-PP2A axis, PKA or another unknown pathway. In agreement with these results, we also observe condition-specific changes in the phosphorylation of proteins involved in rRNA synthesis and translation in the global hierarchical clustering for the different nutrient limitations (Fig 5 B,C).

We mapped phosphorylation sites identified to the MAPK pathway from KEGG (Fig 6). The canonical HOG pathway consists of two MAPK branches converging upon the MAPKK Pbs2, which phosphorylates the MAPK Hog1. Activated Hog1 is imported to the nucleus and mediates the upregulation of ~600 genes via phosphorylation and/or interaction with various transcriptional activators and repressors including Msn2, Msn4, and Sko1 (Westfall, 2004). Here we could not observe a distinct phosphorylation signature in the osmotic stress MAPK cascade for the high salt condition, possibly due to the transient nature of Hog1 activation peaking at 30-60s (Kanshin et al., 2015) and/or the salt induced response overlapping with other stimuli (Vaga et al., 2014). However, we found that phosphorylation of known effectors of the pathway were upregulated after salt stress (Msn2, Msn4, Sko1, Sic1, Hsl1, Hsl7) (Fig 6).

**Figure 6.**
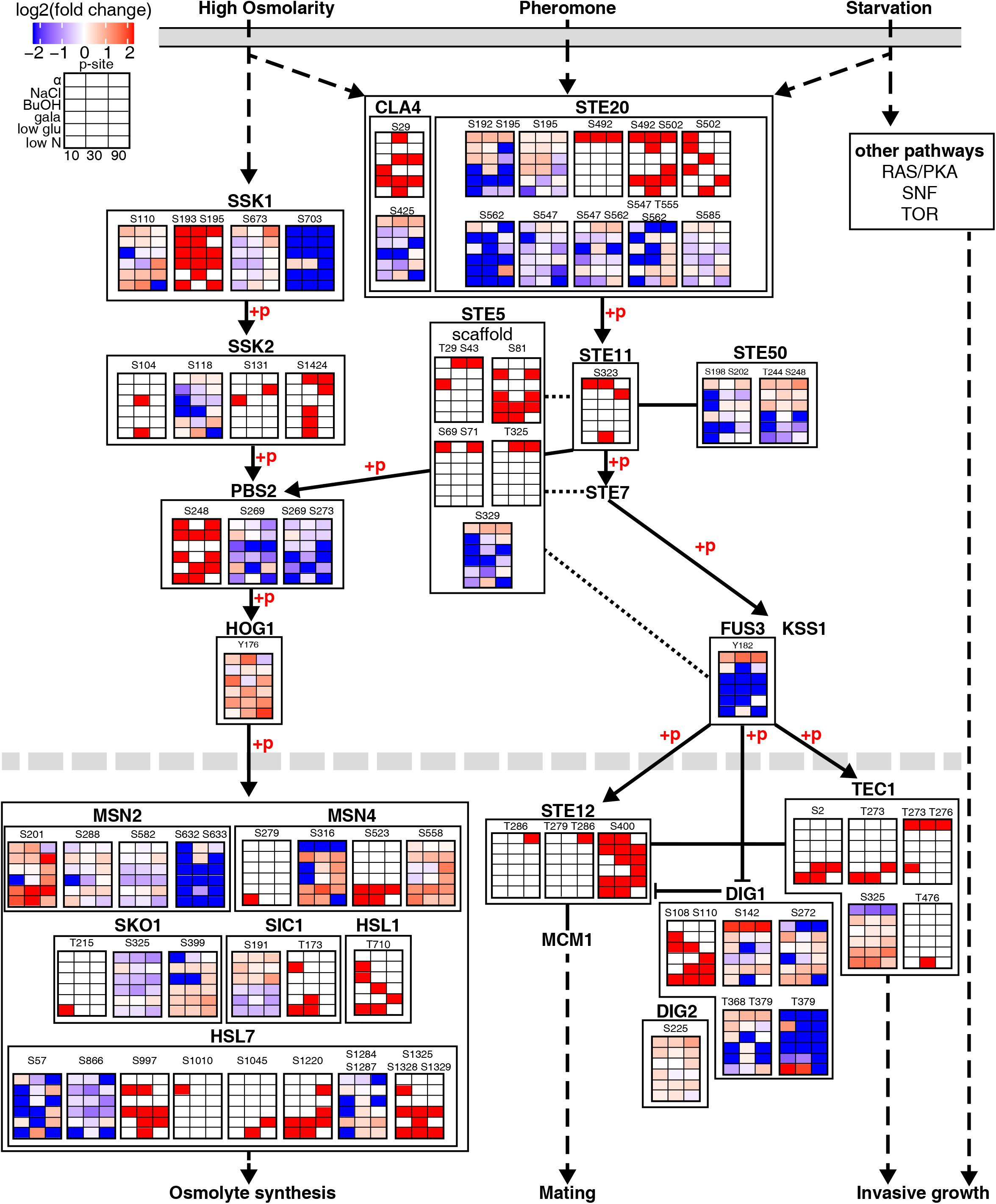
Dynamics of phosphosites in proteins of the canonical MAPK pathways. Representation of partial mating, high osmolarity and invasive growth MAPK KEGG pathway from the plasma membrane to the nucleus. Each heatmap displays log2 fold intensity changes over untreated of one or multiple co-occurring localized phosphorylation site(s) belonging to the indicated proteins. Heatmap rows correspond to treatments and columns to time points. Kinases substrate relationships (+p) and activating or inhibiting effect as stated by KEGG are indicated.

In contrast, with alpha factor stimulation we found stimuli-specific, and often sustained, induction of phosphosites on the kinase Ste20, the pheromone-responsive MAPK scaffold Ste5, the MAPKKK Ste11 and the MAPK Fus3, the adaptor Ste50, and the downstream effectors Ste12, Dig1, Dig2 and Tec1 (Fig 6).

Fus3 activates the mating response and suppresses the invasive growth (Breitkreutz and Tyers, 2002) by phosphorylating and tightly regulating the abundance of transcription factor Tec1 (Breitkreutz et al., 2001; Chou et al., 2004). As expected, we observe co-occurring phosphorylation at T273 and T276, a known phosphodegron (Bao et al., 2010), for all alpha factor stimulation timepoints. However, we also see this Tec1 dual phosphorylation at 10-min glucose limitation and singly phosphorylation of T273 at the early timepoints of nitrogen starvation, indicating that nutrient limitation temporarily suppresses invasive growth. Tec1 activity and stability is regulated on its C-terminus by other not well understood mechanisms (Heise et al., 2010; Köhler et al., 2002) including regulation trough TORC1 (Brückner et al., 2011). We found that dynamic C-terminal phosphorylation at S325 opposes T273/T276 phosphorylation, and we postulate this site could be activating Tec1 by either stabilizing the protein, promoting its nuclear localization, or mediating binding to DNA or to other transcriptional machinery.

Taken together, our multiple perturbation experiment identified informative and robust phosphorylation events on regulatory proteins and transcription factors; as well as effectors where multiple MAPK pathways converge such as Tec1, Msn4 and Stb3. Higher density time course experiments can bring additional resolution to the dynamic response of the MAPK signal transduction cascade, particularly for branches where phosphorylation is transient.

## Discussion

In this work we have established R2-P2, a novel, automated phosphoproteomic sample preparation method that uses magnetic beads for both protein and peptide cleanup and phosphopeptide enrichment. R2-P2 is high-throughput (96-samples in parallel), rapid (1.5 days from cell lysate to proteomic and phosphoproteomic MS injections), robust (compatible with a variety of lysis buffers and detergents), reproducible (coefficient of variation for label-free phosphopeptide quantifications < 25%), high-fidelity (> 90% phosphopeptides purity), high sensitivity (~4,000 phosphopeptides from 25 µg yeast lysate), and low cost for consumables (< $ 3.50 for 250 µg protein). We have shown that R2-P2 is versatile with regards to the phosphopeptide enrichment chemistry, where a combination of different materials can yield deeper coverage of the phosphoproteome. R2-P2 showed comparable or better results than in solution digestion combined with C18 clean up, judged by number of identifications and reproducibility. As a note of caution, protease and phosphatase inhibitors are not efficiently removed by R2-P2 and can lead to significant inhibition of protein digestion or phosphopeptide enrichment, respectively. To facilitate implementation of the R2-P2 protocol in other laboratories, we provide a detailed protocol and a downloadable program for the magnetic bead processor.

Even though the yeast MAPK pathway is one of the best understood signaling systems in biology, our understanding of signal integration and cross talk with other pathways remains superficial. Global and pathway-centric analysis of phosphorylation changes induced by 6 different perturbations at 3 different timepoints reveals a network of interlocking events, rather than a set of simple linear pathways. This is evidenced by both phosphosites regulated under multiple perturbations (phosphosite level crosstalk) and proteins with multiple phosphosites that are differentially regulated (protein level crosstalk). Interestingly, MAPK branches feature different signaling dynamics with regards to their induction and recovery. Pathway-centric analysis recapitulated selective activation of the canonical HOG and mating MAPK pathways. Responses to nutrient starvation were more convoluted, likely due to signal inputs from additional kinases (e.g. Pka and Snf1). With the limited temporal exploration presented we were able to observe robust phosphorylation of transcription factors and regulatory proteins, however the temporal dynamics of the MAPK pathways shall be better recapitulated with additional early timepoints.

The R2-P1 and R2-P2 methods presented here will be attractive for any proteomic laboratory dealing with large sample batches, limited sample material, and challenging biological specimens. We have shown the advantages of R2-P2 for global phosphoproteomic studies, however the method can be extended to automate the enrichment of peptides harboring other post-translational modifications, as far as enrichment chemistries can be added on to magnetic beads. For example, we anticipate future implementations of R2-P2 for peptide immunoaffinity enrichment of acetylation, ubiquitination, tyrosine phosphorylation, and other modifications or motifs. Due to its high sample capacity and robustness, R2-P2 can become a cornerstone method to study protein phosphorylation and regulatory signaling networks in biology and disease.

## Methods

### Cell culture and treatment

For method optimization experiments we used *S. cerevisiae* strain BY4741 and for stimulation experiments strain Σ1278b MATa. Yeast were cultured overnight at 30°C in synthetic complete medium (SMC) with 2% glucose. Cells were subsequently inoculated at OD_600_ = 0.1 and grown to OD_600_ = 0.6. Cells were harvested by adding 100% trichloroacetic acid (w/v) to a final concentration of 10% trichloroacetic acid, incubated on ice for 10 min, centrifuged, decanted, washed once with 100% ice-cold acetone, centrifuged, decanted, snap frozen and pellets were stored at −80°C.

Following media were used to treat the Σ1278b strain: 20 mM alpha factor in SMC-2% glucose, SMC-0.2% glucose, 0.4 M NaCl in SMC-2% glucose, 1% 1-butanol in SMC-2% glucose, SMC-2% galactose and SLAD - low nitrogen media (1.7% YNB, 50 mM Ammonium sulfate, 2% glucose, leucine, uracil, tryptophan, histidine). To treat Σ1278b, cells were grown to OD_600_ = 0.6, harvested by centrifugation, washed twice with prewarmed new media base (SMC-2% glucose, SMC-0.2% glucose, SMC-2% galactose or SLAD) and then treatments were induced for 10 min, 30 min and 90 min followed by trichloroacetic acid harvest as described above.

### Cell lysis, protein reduction, and alkylation

Frozen cell pellets were resuspended in lysis buffer composed of 8 M urea, 75 mM NaCl and 100 mM Tris pH 8.2. Cells were lysed by 3 rounds of bead beating (30 s beating, 1 min rest) with zirconia/silica beads. Lysate protein concentration was measured by BCA assay. Proteins were reduced with 5 mM dithiothreitol (DTT) for 30 min at 55°C and alkylated with 15 mM iodoacetamide in the dark for 15 min at room temperature. The alkylation reaction was quenched by incubating with additional 5 mM DTT for 15 min at room temperature.

### Paramagnetic beads

For R2-P2 a 1:1 mix of carboxylated paramagnetic beads (Sera-Mag SpeedBeads, CAT# 09-981-121, 09-981-123) were used. Beads come at a stock concentration of 50 µg/µl. Before usage the two different types of beads were combined, washed 3 times with water and diluted to the working concentration of 10 µg/µl in water.

For phosphopeptide enrichments the following paramagnetic beads were used: PureCube Fe^3+^-NTA MagBeads (Cube Biotech), Ti^4+^-IMAC (MagReSyn), Zr^4+^-IMAC (MagReSyn) and TiO_2_ (MagReSyn). Paramagnetic beads from MagReSyn were used as provided and equilibrated according to the manufacturer protocol. If not stated otherwise, Fe^3+^-NTA MagBeads were used.

### R2-P1 protocol

The R2-P1 purification and digestion was implemented in the following way on a KingFisher™ Flex (Thermo Scientific): The 96-well comb is stored in plate #1, magnetic carboxylated beads (1 µl of 10µg/µl carboxylated bead mix per 1 µg protein) in plate #2, lysate-ethanol mixture (0.5 µg/µl protein, 50% ethanol v/v) in plate #3, wash solutions (same volume as lysate-ethanol mixture, 80% ethanol) in plates #4 to #6, elution/digestion buffer (>50 µl, 25 mM ammonium bicarbonate, pH 8) together with digestion enzyme in plate #7 and second elution (water) in plate #8 (Fig 1A). According to the required volumes shallow 96-well kingfisher plates (50 µl - 150 µl) or deep 96-well kingfisher plates (150 µl - 1000 µl) were chosen.

A program was compiled to collect the carboxylated beads in plate #2, move them to plate #3 for protein binding and subsequently to plate #4, #5 and #6 for protein desalting and purification. The protein purification takes 30 minutes and the protocol pauses to allow for loading of plate #7 containing the digestion enzyme, which allows for preparation of the digestion solution only immediately before use. The beads are then moved to plate #7 and proteins are eluted/digested at 37 ºC for 3.5 hours with constant agitation. Carboxylated beads are subsequently moved and washed in the second elution plate #8 and afterwards discarded. At this step the robotic program ends and plates can be removed. Elution in plate 1 and 2 are combined in one plate, acidified with formic acid and clarified by centrifugation. At this point aliquots are taken for total proteome analysis. For this, the required amount of peptides is transferred to an MS sample vial plate or individual vials and dried down. The total proteome samples can be stored or resuspended in MS loading buffer and measured by LC-MS/MS. Optional, but recommended when processing a lot of samples in parallel: to avoid any carryover of beads into the LC system, sample can be filtered through a C8 stage tip prior to drying. The lyophilized peptides in the 96-well plate are resuspended in phosphopeptide enrichment binding buffer and automated phosphopeptide enrichment is performed for R2-P2.

### R2-P2 protocol and automated phosphopeptide enrichments

The automated phosphopeptide enrichment was implemented in the following way on the KingFisher™ Flex (Thermo Scientific): The 96-well comb is stored in plate #1, magnetic Fe^3+^-IMAC, Ti^4+^-IMAC, Zr^4+^-IMAC, or TiO_2_ beads in plate #2, resuspended peptides in plate #3, wash solutions in plates #4 to #6 and elution solution in plate #7 (Fig 1A). Shallow 96-well kingfisher plates were used.

A program was compiled to collect the magnetic beads in plate #2, move them to plate #3 for phosphopeptide binding and subsequently to plate #4, #5 and #6 for washing. The phosphopeptide purification takes 40 minutes and the protocol pauses to allow for loading of plate #7 containing the elution solution, which allows for preparation of the elution solution immediately before use, avoiding evaporation of ammonia. The beads are subsequently moved to plate #7 and phosphopeptides are eluted. Plates are removed from the robot at this point and the elution is immediately neutralized by adding 30 µl 10% formic acid in 75% acetonitrile to each well. For the Fe^3+^-IMAC protocol binding and wash solutions consisted of 150 ul 80% acetonitrile and 0.1% trifluoroacetic acid and elution solution was 50 µl 50% acetonitrile and 2.5% NH_4_OH. For Ti^4+^-IMAC, Zr^4+^-IMAC or TiO_2_ buffer composition were chosen according to the recommendation of MagReSyn. After acidification of the elution, peptides were dried down in a speed-vac stored at −20 ºC or resuspended in MS loading buffer and analyzed by LC-MS/MS. Optional but recommended when processing a lot of samples in parallel: to avoid any carryover of beads into the LC system, sample can be filtered through a C8 stage tip prior to drying.

### Mass spectrometry data acquisition

Dried peptide and phosphopeptide samples were dissolved in 4% formic acid, 3% acetonitrile and analyzed by nLC-MS/MS. Peptides were loaded onto a 100 µm ID x 3 cm precolumn packed with Reprosil C18 3 µm beads (Dr. Maisch GmbH), and separated by reverse phase chromatography on a 100 µm ID x 30 cm analytical column packed with Reprosil C18 1.9 µm beads (Dr. Maisch GmbH) and housed into a column heater set at 50ºC.

Method optimization experiments were performed on a Q Exactive Hybrid Quadrupole-Orbitrap mass spectrometer (Thermo Fisher) equipped with an Easy1000 nanoLC (Thermo Fisher) system. Phosphopeptides were separated by a gradient ranging from 7 to 28% acetonitrile in 0.125% formic acid over 60 min. For DDA experiments full MS scans were acquired from 350 to 1500 m/z at 70,000 resolution with fill target of 3E6 ions and maximum injection time of 100 ms. The 20 most abundant ions on the full MS scan were selected for fragmentation using 2 m/z precursor isolation window and beam-type collisional-activation dissociation (HCD) with 26% normalized collision energy. MS/MS spectra were collected at 17,500 resolution with fill target of 1E5 ions and maximum injection time of 50 ms. Fragmented precursors were dynamically excluded from selection for 30 s.

DDA for spectral library generation, DIA for gas-phase fractionation, and DIA for quantitation were performed on an Orbitrap Fusion Lumos Tribrid Mass Spectrometer (Thermo Fisher) equipped with a nanoACQUITY nanoLC (Waters). Phosphopeptides were separated by a gradient ranging from 7 to 28% acetonitrile in 0.125% formic acid over 60 min. All MS spectra were acquired on the orbitrap mass analyzer and stored in centroid mode. For the DDA experiments full MS scans were acquired from 350 to 1500 m/z at 60,000 resolution with an AGC target of 7e5 ions and maximum injection time of 50 ms. The most abundant ions on the full MS scan were selected for fragmentation using 1.6 m/z precursor isolation window and beam-type collisional-activation dissociation (HCD) with 28% normalized collision energy for a cycle time of 3s. MS/MS spectra were collected at 15,000 resolution with fill target of 5e4 ions and maximum injection time of 50 ms. Fragmented precursors were dynamically excluded from selection for 40 s. For the DIA gas-phase fractionation, the Fusion Lumos was configured to acquire seven runs, each spanning ~100 m/z (448.475-552.524, 548.525-650.574, 648.575-750.624, 748.625-850.674, 848.675-950.724, 948.725-1050.774, 1048.775-1150.824). Full MS scans were acquired in centroid mode from 440 to 1160 m/z at 60,000 resolution, AGC target of 1e6 and maximum injection time of 100ms. MS/MS were acquired at 30,000 resolution, using 4 m/z precursor isolation window, an AGC target of 2e5 and maximum injection time of 54 ms, and HCD with 28% normalized collision energy. For quantitative DIA analysis, the Fusion Lumos was configured to acquire a full MS scan in centroid mode from 350 to 1500 m/z at 60,000 resolution, AGC target 7e5 and maximum injection time 50 ms followed by 48 × 15 m/z DIA spectra using 1 m/z overlapping windows from 472.489 to 1,145.489 m/z at a 15,000 resolution, AGC target 5e4 and maximum injection time of 22 ms.

### Mass spectrometry data analysis

DDA MS/MS spectra were searched with Comet (Eng et al., 2013) against the *S. cerevisiae* proteome. The precursor mass tolerance was set to 20 ppm. Constant modification of cysteine carbamidomethylation (57.021463 Da) and variable modification of methionine oxidation (15.994914 Da) were used for all searches and variable modification of serine, threonine, and tyrosine phosphorylation (79.966331 Da) was used for phosphopeptide samples. Search results were filtered to a 1% FDR at PSM level using Percolator (Käll et al., 2007). Phosphorylation sites were localized using an in-house implementation of the Ascore algorithm (Beausoleil et al., 2006). Phosphorylation sites with an Ascore > 13 (P < 0.05) were considered confidently localized. Peptides were quantified using in□house software measuring chromatographic peak maximum intensities.

For overlapping DIA runs MSConvert (Chambers et al., 2012) was used to deconvolute RAW files and generate mzMLs. A spectrum library was created from the Fusion Lumos DDA data using Skyline (version 4.2.0.19009) (MacLean et al., 2010). This BLIB library was imported to EncyclopeDIA (version 0.8.1) (Searle et al., 2018b), used to search the DIA gas-phase fractionated runs and a chromatogram library was generated. This ELIB file was imported to Thesaurus (version 0.6.4) (Searle et al., 2018a) and quantitative DIA files were searched using a percolator FDR threshold > 0.01 and the option “recalibrated (peak width only)” was chosen as a localization strategy.

### Bioinformatic analysis

Bioinformatic analysis was performed using R (https://www.r-project.org/) and Perseus (Tyanova et al., 2016). Quantitative values obtained from DDA and DIA analysis were median normalized.

For all boxplots the lower and the upper hinges of the boxes correspond to the 25% and the 75% percentile, and the bar in the box the median. The upper and lower whiskers extend from the largest and lowest values, respectively, but no further than 1.5 times the IQR from the hinge.

For the DIA experiments phosphopeptides were only quantified if they were observed in at least 2 out of 3 biological replicates. After filtering, missing values were globally imputed according to the background. For phosphosite centric analyses only phosphopeptides were considered where the site could be localized based on site-specific fragment ions at least once and no conflicting positional isomers were localized in other samples.

All correlation calculations utilize the Pearson method. The calculation of individual p values was performed using two sided Student’s t tests of biological replicates, unless otherwise noted. To obtain data for hierarchical clustering and heatmap plots, ANOVA multiple-sample tests were performed on intensity values of phosphopeptides and data was filtered for values with Benjamini–Hochberg multiple-hypotheses corrected q-value of <0.05. For hierarchical clustering Z-scored values were used as input (Fig 5B) and for all other heatmaps log2 fold changes over the untreated were used.

Principal component analysis (PCA) and term enrichment analyses were performed using Perseus. Term enrichment was performed, by annotating phosphopeptides with the corresponding protein annotation terms, and performing Fisher exact testing on the level of the protein. The categories used for annotation of proteins were Gene Ontology (GO) Biological Processes slim, GO Molecular Functions and Kyoto Encyclopedia of Genes and Genomes (KEGG). Observed differences were filtered for a Benjamini–Hochberg multiple-hypotheses corrected q-value of <0.02 for enrichments in hierarchical clustering and q <0.05 for category enrichment in components of the PCA.

### Data availability

All mass spectrometry proteomics data have been deposited to the ProteomeXchange Consortium via the PRIDE partner repository (Perez-Riverol et al., 2019) with the dataset identifier PXD013453. Reviewer account details: Username: Password: nx7792Pu

## Author contributions

M.L., R.R.-M., and J.V. conceived the study. M.L. conducted most of the experiments, with assistance from N.L. and advice from R.R.-M. M.L. analyzed the data. J.V. supervised the study. M.L. and J.V. wrote the paper and all the authors edited it.

## Acknowledgments

We thank Ian Smith for advice with the MS measurements, Anthony Valente for bioinformatic support, and all members of the Villén lab for useful discussions. M.L. is supported by a postdoctoral fellowship from the Swiss National Science Foundation (P2ZHP3_181503). This work is supported by NIH grants R35GM119536 and R01AG056359, Human Frontier Science Program grant RGP0034/2018, a Research program grant by the W.M. Keck Foundation, and the University of Washington’s Proteomics Resource UWPR95794.

---

**Supplementary Figure S1.**
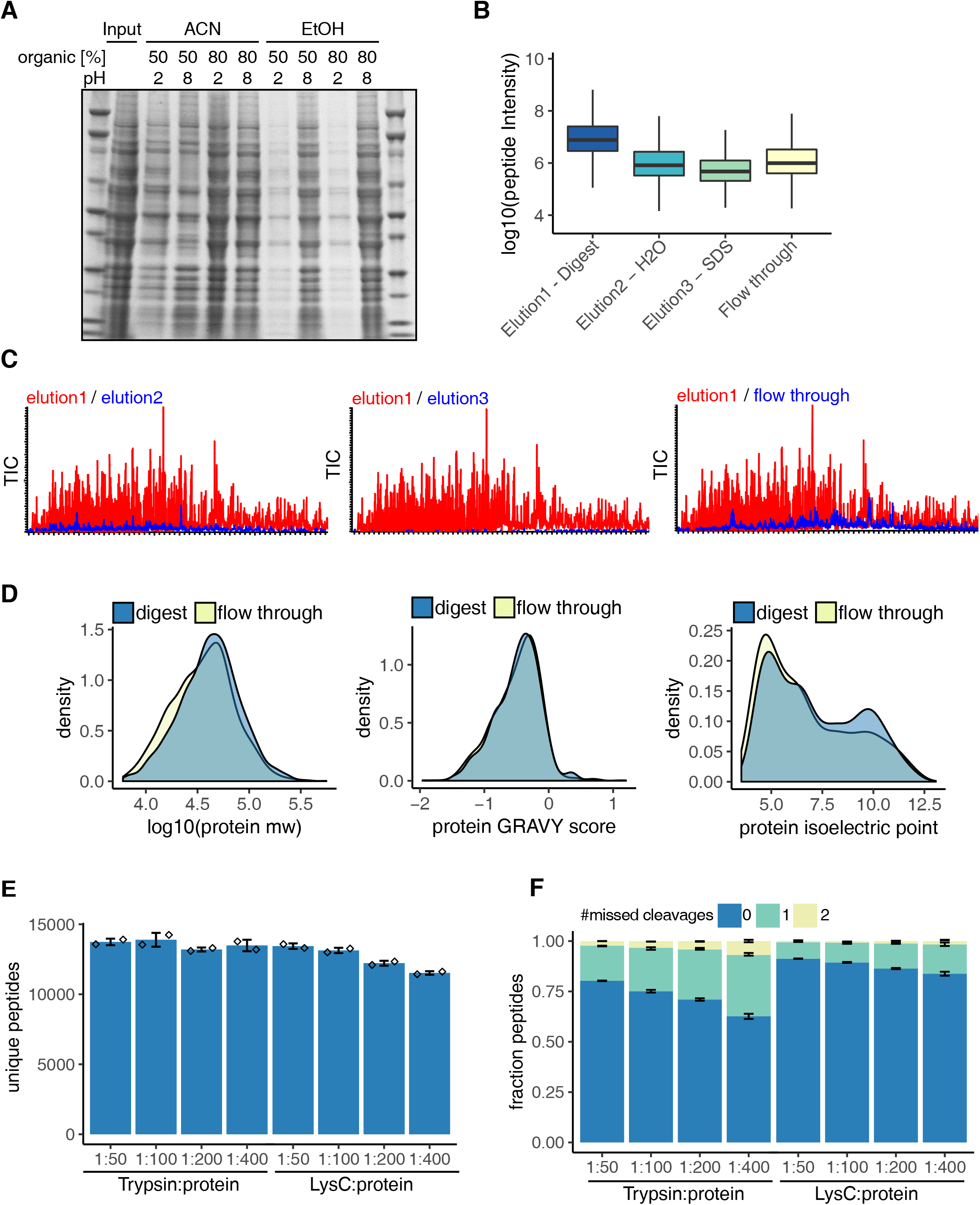
Evaluation of protein capture, desalting, digestion and elution efficiency for R2-P1. A) Binding and washing conditions for R2-P1 were tested on 50 µg yeast lysate. Yeast lysate were brought to pH 2 or pH 8 respectively and ethanol or acetonitrile was added to reach the indicated dilutions (v/v). Beads were washed 3 times in the corresponding binding buffer. Proteins were eluted and resolved on a SDS-PAGE. B) 50 µg of yeast protein extract were processed and digested using R2-P1. Elution 1 consisted of 25mM ammonium bicarbonate with trypsin, elution 2 was water and elution 3 was 1% SDS. The flow through from the magnetic beads binding step was dried and digested in solution with trypsin. All 4 samples were additionally desalted using C18 stage tips in this case. 1/50 of each sample were injected on the MS. Shown is a boxplot of the MS1 peptide intensity measured for the different samples. C) Total ion currents (TIC) of the samples measured in B). D) Density plots of the distribution of molecular weight, GRAVY score and isoelectric point of the proteins identified in the R2-P1 processed samples and the flow through of the R2-P1 protein binding step. E) Samples consisting of 50 µg yeast protein extract were prepared by R2-P1 and digested using trypsin or LysC at an enzyme:protein ratio of 1:50, 1:100, 1:200 and 1:400 respectively. Mean peptide IDs are displayed (n=2) F) Fraction of missed cleaved peptides from E).

**Supplementary Figure S2.**
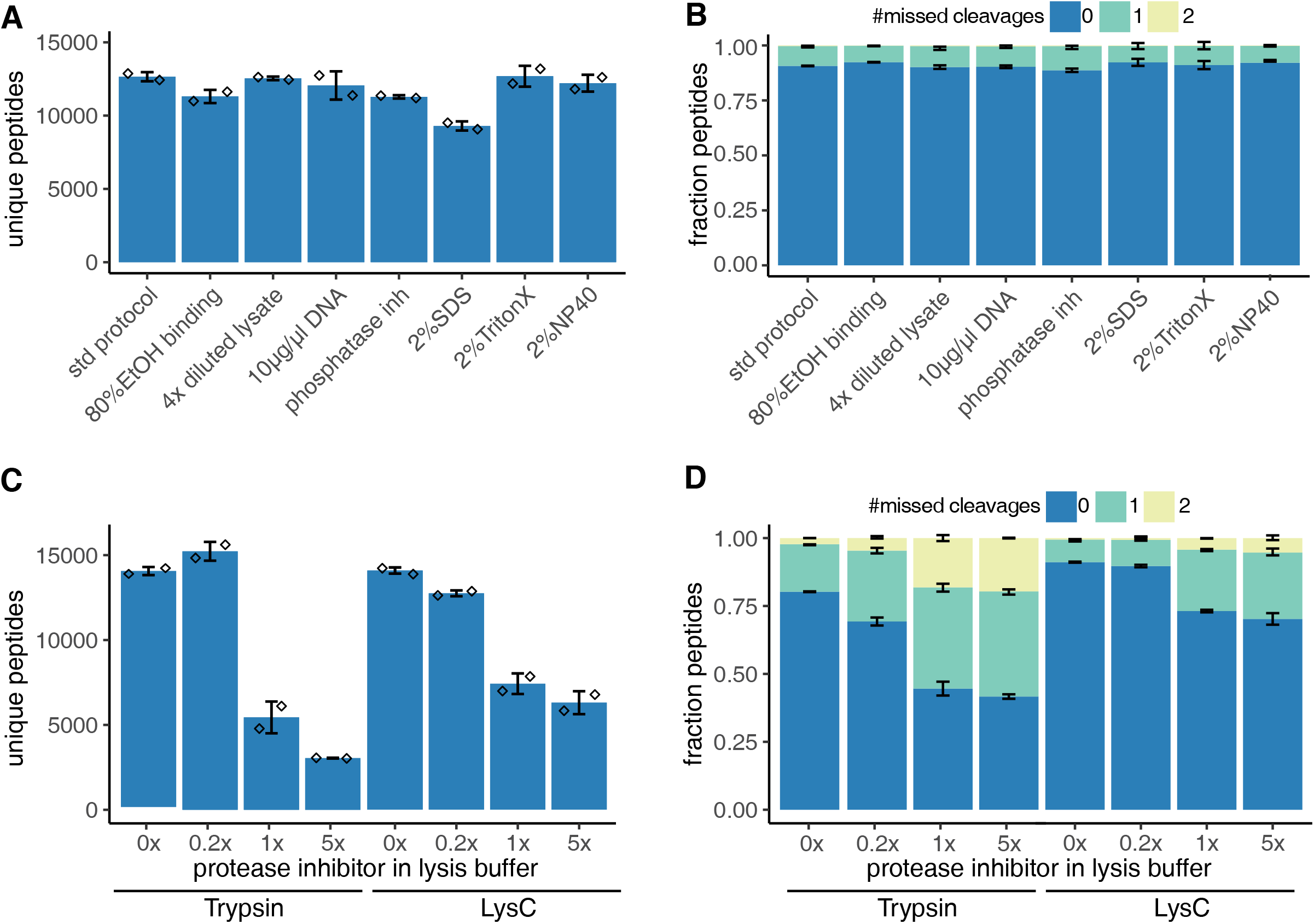
Evaluation of lysis buffer composition for R2-P1 for total proteome analysis. 200µg of yeast protein extract in different buffer compositions were processed by R2-P1 followed by peptide analysis with DDA-MS. A) Mean count of unique peptides are shown for the different conditions (n=2). B) Fraction of peptides with 0, 1 or 2 missed cleavage sites identified in A). The mean is displayed (n=2). C) Different amounts of protease inhibitor (Pierce) was added to 50 µg yeast protein extract to reach 0.2x, 1x and 5x protease working concentrations relative to the binding and digestion buffer volume. Samples were processed by R2-P1 using either trypsin or lysC for digestion. Mean peptide IDs are displayed (n=2). D) Fraction of phosphopeptides with 0, 1 or 2 missed cleavage sites identified in C). The mean is displayed (n=2).

**Supplementary Figure S3.**
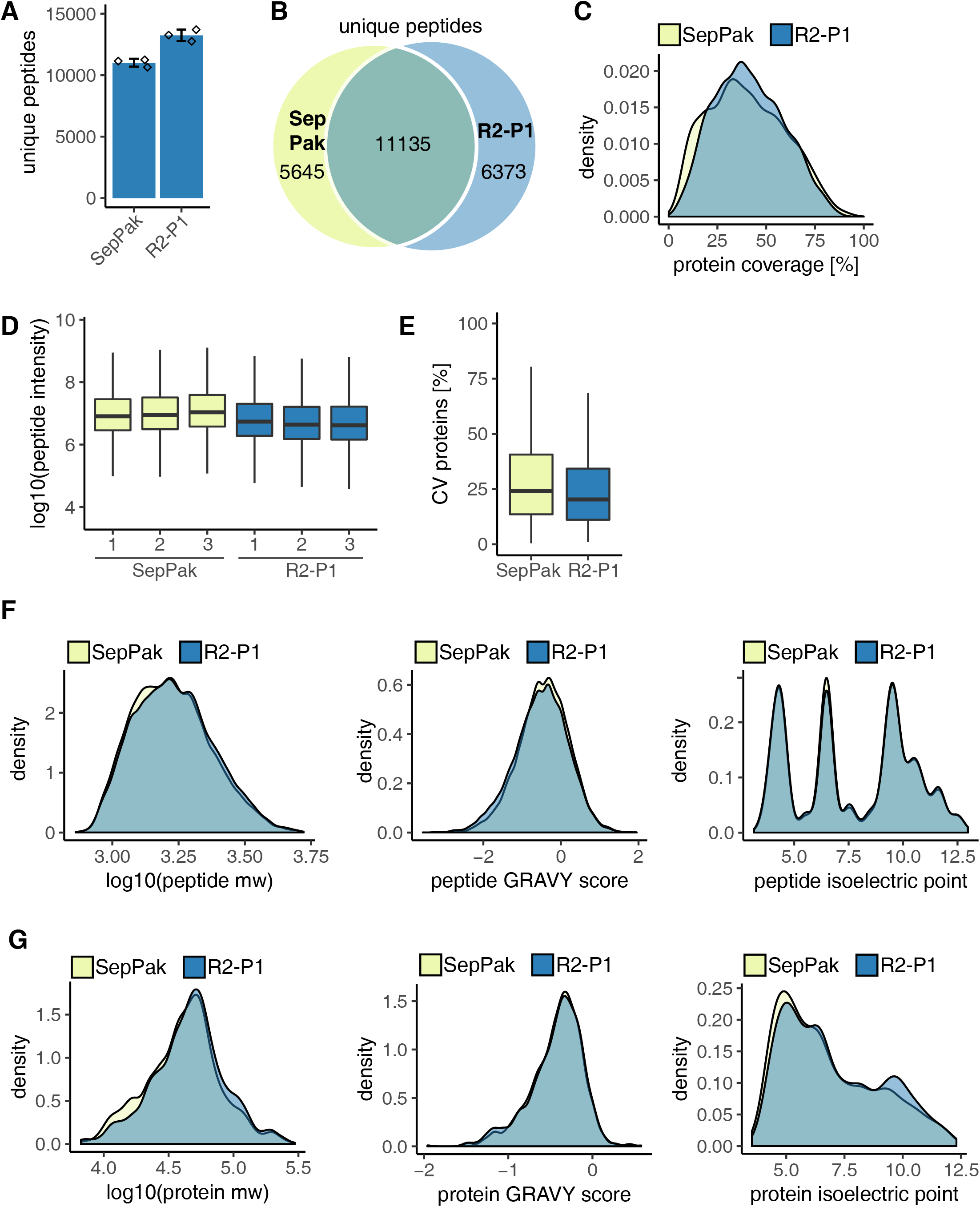
Comparison of total proteome analysis with R2-P1 versus in solution digestion C18 SepPak desalting. 25 µg of yeast protein extract were processed by R2-P1 or by manual in-solution digestion and reversed phase C18 SepPak desalting in triplicates. Peptide analysis was performed by DDA-MS. A) Mean count of unique peptides are shown (n=3). B) Venn diagram of peptides identified in samples processed by R2-P1 or SepPak. C) Density plot of protein coverage distribution for the two different methods. D) Boxplot of MS1 peptide intensities (n=3). E) CVs of protein intensities for the different sample processing protocols (n=3). F) Density plots of the distribution of molecular weight, GRAVY score and isoelectric point of the proteins identified in the two different sample processing protocols. G) Density plots of the distribution of molecular weight, GRAVY score and isoelectric point of the peptides identified in the two different sample processing protocols.

**Supplementary Figure S4.**
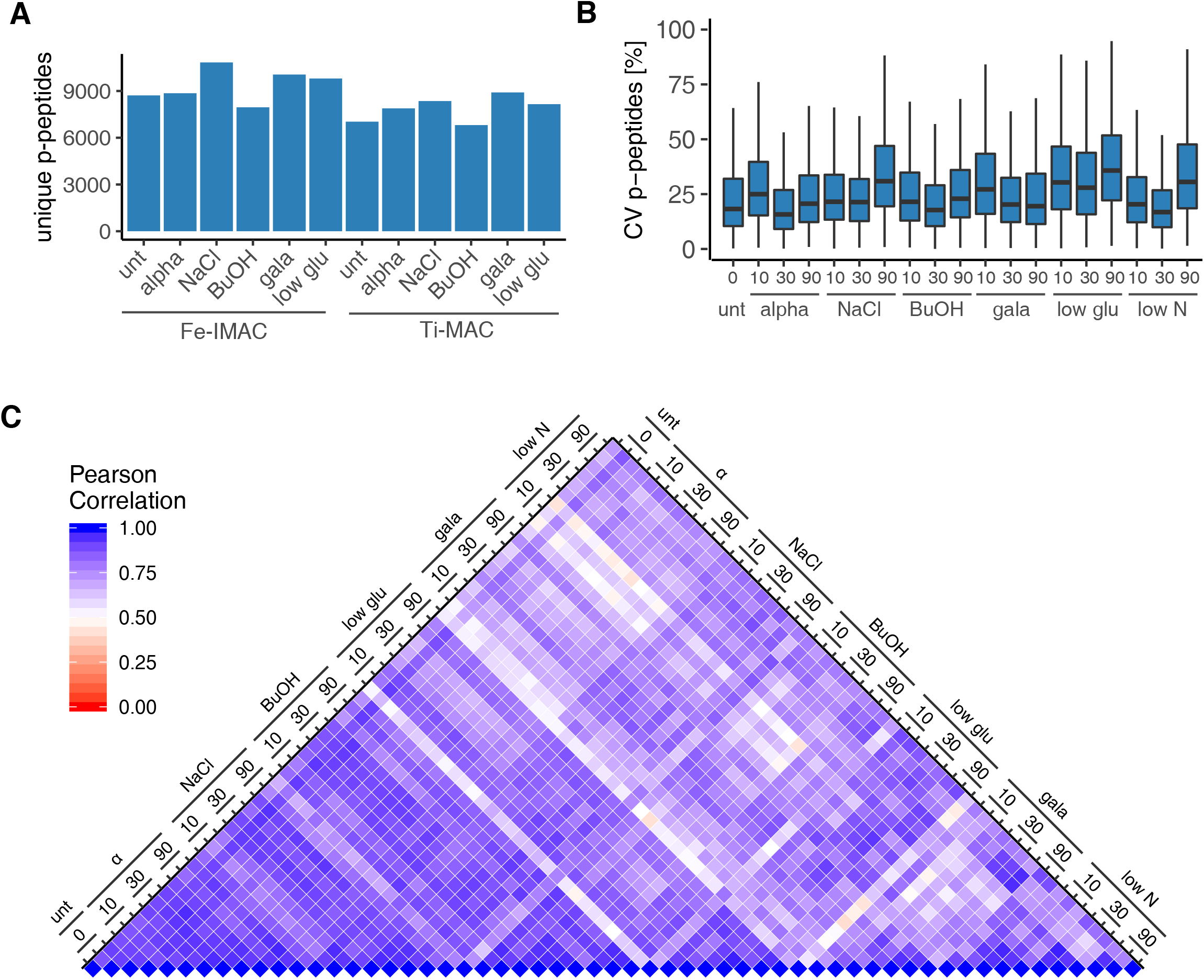
Library generation and R2-P2 DIA-MS reproducibility. A) Identified phosphopeptides in Fe^3+^-IMAC and Ti^4+^-IMAC fractions of pooled time points for the different treatments measured in single injection DDA-MS experiments. B) Coefficient of variation for phosphopeptide quantification in the DIA-MS experiment (n=3). C) Heatmap of Pearson correlation coefficients for all DIA-MS measurements.

**Supplementary Figure S5.**
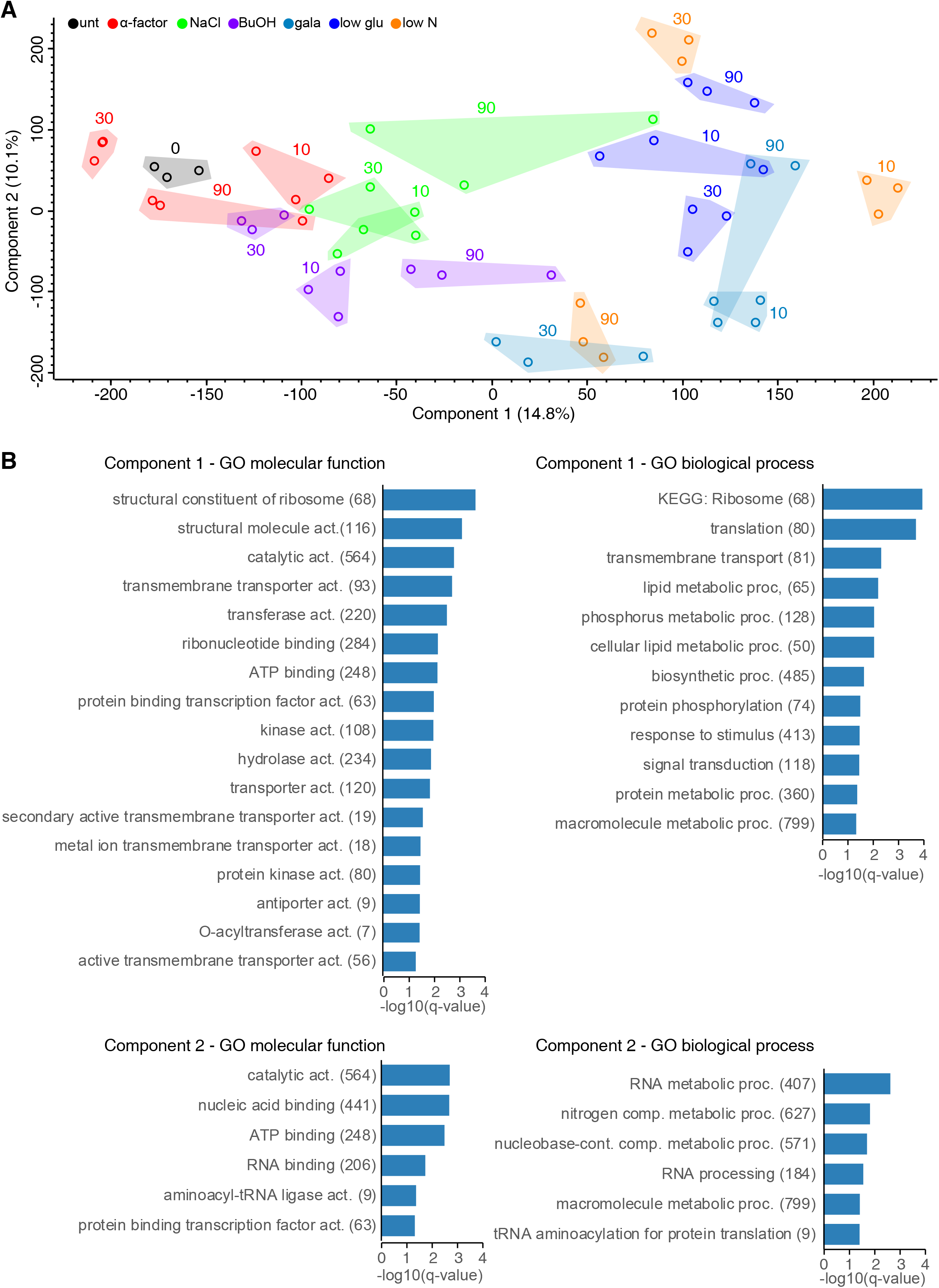
Principal component analysis of all DIA-MS measurements. A) Principal component analysis (PCA) of all phosphoproteomic data for the different stimuli and time points. Color coding depicts different treatments and corresponding time points are indicated in the plot. B) Term-enrichment analysis displaying gene ontology molecular function and biological processes terms that were significantly enriched (Fisher’s exact test with Benjamini– Hochberg multiple-hypotheses correction at q < 0.02) for phosphorylated proteins in component 1 and 2 and their corresponding q-values.

**Supplementary Figure S6.**
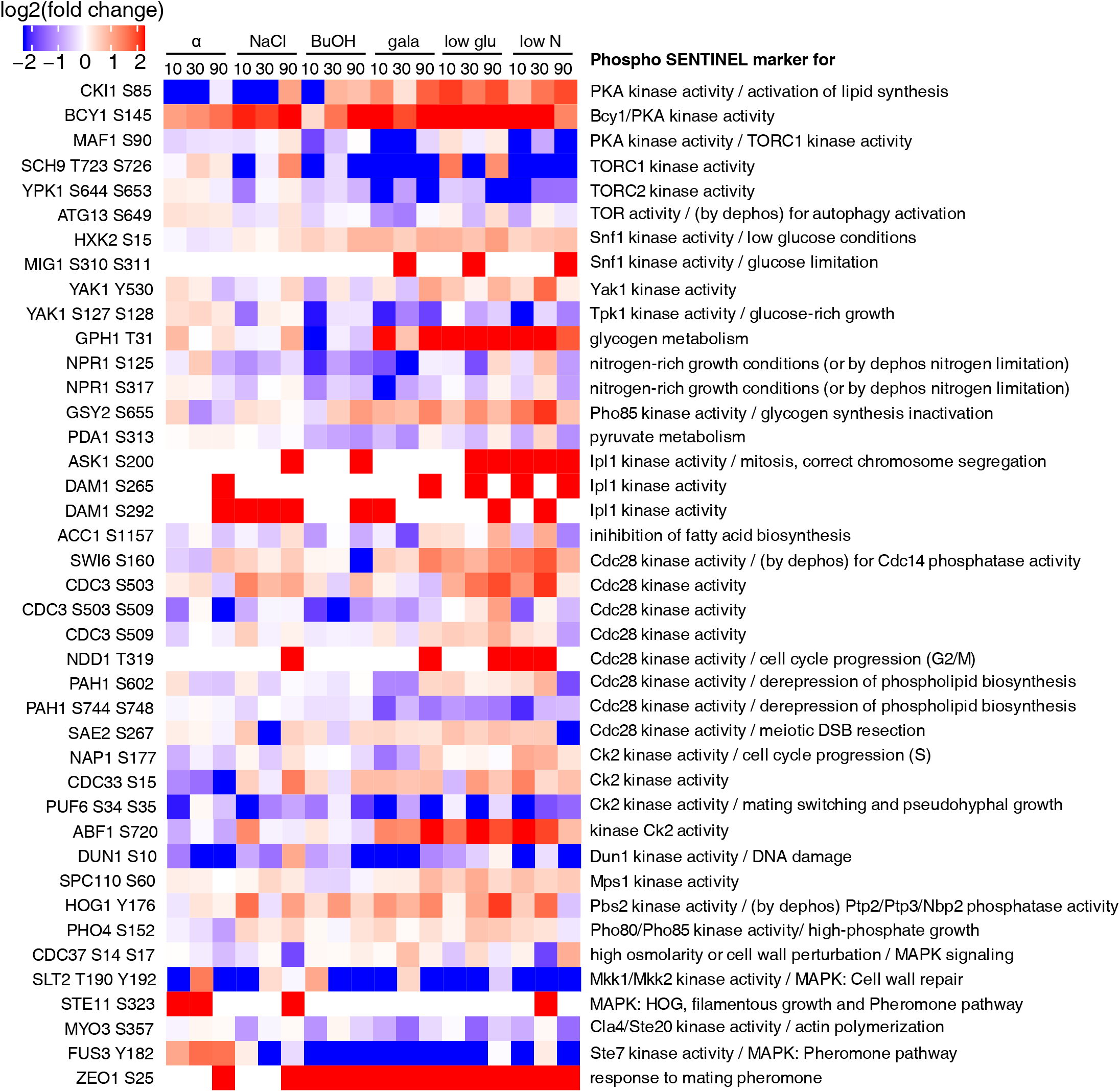
Phosphosite markers for biological processes. Phosphosites were mapped on phospho sentinel marker peptides as defined by Soste et al (Soste et al., 2014). Log2 fold changes over untreated are shown for the different treatment and time points. Phosphosite is indicated on the left side of the heatmap and description for upregulation of the phospho marker on the right.

**Supplementary Figure S7.**
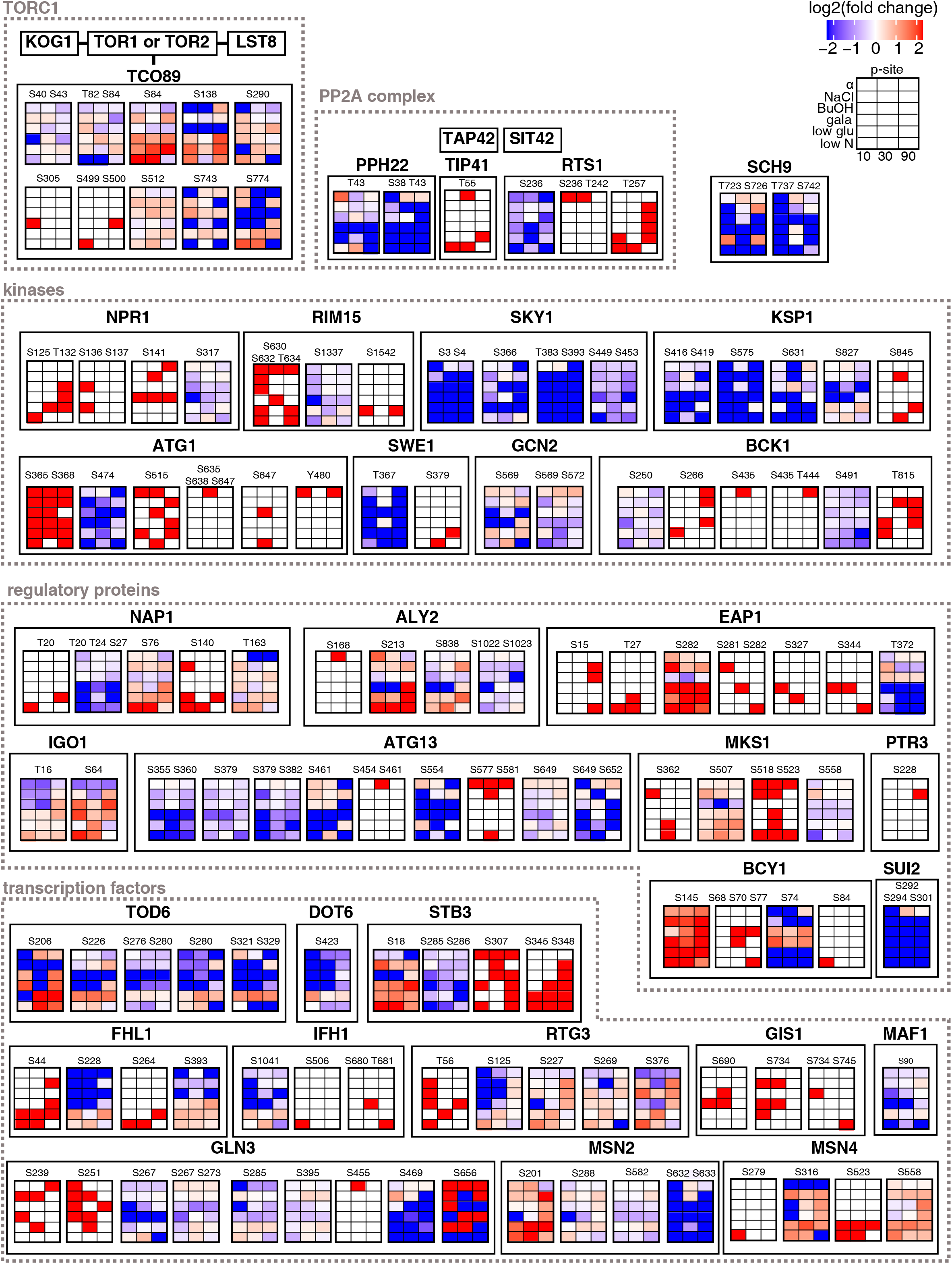
Dynamics of phosphosites for TORC1-related proteins. Quantified phosphosites were mapped onto literature curated TOR targets as reported by Oliveira et al. (Oliveira et al., 2015). Each heatmap displays log2 fold intensity changes over untreated of one or multiple co-occurring localized phosphorylation site(s) belonging to the indicated proteins. Heatmap rows correspond to treatments and columns to time points.

